# Tubulin acetylation governs organelle remodeling and lysosomal reformation during neuronal differentiation

**DOI:** 10.64898/2026.02.13.705749

**Authors:** Chih-Hsuan Hsu, Alexander Josiah Kinrade, Maria Clara Zanellati, Sarah Cohen

## Abstract

A functional nervous system depends on neuronal morphology established during differentiation. The microtubule (MT) cytoskeleton supports neuronal differentiation by organizing organelle positioning and facilitating transport. The dynamics and properties of MTs are regulated by a variety of post-translational modifications (PTMs), with many organelle interactions occurring preferentially on modified MTs. Here we find that tubulin acetylation is enriched at specific subcellular locations during differentiation of human induced neurons. We apply a quantitative multispectral imaging pipeline to simultaneously analyze eight membrane-bound organelles and define how tubulin acetylation reshapes organelle architecture and interaction networks during neuronal differentiation. We find that loss of tubulin acetylation broadly alters organelle morphology, spatial distribution, and inter-organelle interactions, with lysosome-organelle interactions most affected. Loss of acetylated MTs leads to enlarged, highly acidified lysosomes, impaired lysosomal fission, and accumulation of autolysosomes, consistent with defective lysosomal reformation. Super-resolution microscopy further reveals that lysosome-endoplasmic reticulum (ER) contacts preferentially associate with acetylated MTs. Together, our data support a model in which tubulin acetylation coordinates lysosome-ER interactions to facilitate lysosome remodeling and turnover. This work establishes tubulin acetylation as a key cytoskeletal regulator that links organelle interactions to organelle homeostasis important for neuronal differentiation.

## Introduction

Neuronal differentiation is a highly coordinated process that converts morphologically simple stem cells into neurons, one of the most structurally and functionally complex human cell types. This process requires extensive cytoskeletal, membrane, and organelle remodeling to set up the functional units of the nervous system^1,2^. As differentiation occurs, neural stem cells undergo morphogenesis to develop their polarity and elaborate functionally specialized compartments, axons and dendrites^3,4^. It is well established that microtubules (MTs) are critical cytoskeletal elements in neurons for structural support, building neuronal morphology, positioning membrane-bound organelles^5^ and serving as railroads for long-distance organelle transport^6–8^. The dynamics and properties of MTs are regulated by different post-translational modifications (PTMs), the “tubulin code”^9,10^. Among tubulin PTMs, the acetylation of lysine 40 (K40) on α-tubulin is unique for protecting long-lived MTs from mechanical stress by enhancing their flexibility and resistance^11,12^. Multiple studies demonstrated that tubulin acetylation is central for proper neuron development, where it regulates neuron migration^13^, morphological transition^14,15^ and dentate gyrus formation^16^.

In parallel, studies have shown that metabolic transition from glycolysis to oxidative phosphorylation happens during neuronal differentiation accompanied by massive multi-organelle rearrangements which are essential for generating functional neurons^17,18^. For example, temporal proteomics revealed that endoplasmic reticulum (ER), Golgi, lysosome and mitochondria are substantially remodeled by canonical autophagy^19^. ER undergoes selective ER-phagy^20^, somatic Golgi translocates into neurites^21^, lysosomes shrink and become more motile^22^ and mitochondria elongate along with the metabolic switch to oxidative phosphorylation^17,23,24^. These examples highlight the contribution of individual organelles to neuronal differentiation. However, organelles do not function independently. They communicate through membrane contact sites, where the coordinated signaling and the exchange of ions, lipids, metabolites, and proteins take place^25^. Beyond organelle morphology and distribution, inter-organelle contact sites are integral to cell functionality^26–28^ and are also actively remodeled during neuronal differentiation^18,29,30^.

Tubulin acetylation regulates multiple organelle processes in cultured non-neuronal cells. For example, lysosomes move on perinuclear acetylated tracks by kinesin-1, ER tubule extension is found along acetylated MTs for ER-mitochondrial contacts and the Golgi is embedded within the network of crosslinked acetylated MTs^31–33^. In neurons, tubulin acetylation regulates multiple organelle processes including ER tubule extension^34^, as well as mitochondrial and lysosomal transport^35–37^. Although tubulin acetylation has been implicated in neuronal development and in regulating selected organelle processes, it remains incompletely understood whether tubulin acetylation coordinates a broader, multi-organelle remodeling during neuronal differentiation and how changes in acetylated MTs affect the global landscape of organelle architecture including morphology, their interaction networks and spatial distribution at the whole-cell level.

To address this gap, we used a standardized human induced pluripotent stem cell (iPSC)-derived neuron (iNeuron) system to model human neuronal differentiation. iNeurons provide developmentally synchronized, genetically stable and highly reproducible cortical neurons^38^. Using this differentiation platform, we discovered that multiple tubulin PTMs including tubulin acetylation increase as neurons differentiate, with MT acetylation particularly enriched in specific subcellular domains within the soma. To test the role of acetylated MTs in organelle morphology and interactions during neuronal differentiation, we depleted tubulin acetylation by knocking down its primary catalytic enzyme, α-tubulin acetyltransferase (ATAT1). Multispectral confocal microscopy has recently emerged as a powerful tool utilized by our group and others for high-throughput multi-organelle analysis^18,39–42^. We previously identified robust cell type-specific organelle signatures of primary neurons and astrocytes^43^. Therefore, we applied eight-color multispectral imaging to ATAT1 knockdown iNeurons to systematically study the effects of α-tubulin K40 acetylation on organelle organization during neuronal differentiation. We discovered that tubulin acetylation affects the abundance, morphology, and interactions among ER, mitochondria and lysosomes. Loss of tubulin acetylation in iNeurons leads to broad reduction of organelle interactions, with the most prominent decreases in lysosome-ER contacts in both soma and neurites. Super-resolution microscopy further revealed that lysosome-ER contacts display strong spatial association with acetylated MTs. In addition to morphological changes, we observed that tubulin acetylation governs lysosome dynamics by supporting normal fission frequency. Functionally, we observed progressively enlarged and increased acidified lysosomes following prolonged depletion of acetylated tubulin, accompanied by accumulation of autolysosomes, suggesting a defect in lysosome reformation. Overall, our studies establish a platform to dissect the impact of tubulin acetylation on neuronal differentiation at the level of multi-organelle remodeling and reveal a critical role for MT acetylation in sustaining lysosome reformation.

## Results

### Tubulin PTMs are enriched in hiPSC-derived cortical neurons during differentiation

Following an established human iPSC to iNeuron differentiation protocol based on Neurogenin-2 (NGN2) expression^38,44^, we generated a NGN2-KOLF2.1J iPSC stable cell line that can be efficiently differentiated into human neurons^18^, which provides a reliable model to study cytoskeletal organization during neuronal differentiation (**Figure 1A**). Tubulin post-translational modifications (PTMs) play critical roles in regulating microtubule (MT) dynamics^45^, controlling neuronal development^46,47^ and are catalyzed by distinct sets of enzymes (**Figure 1B**). To determine how tubulin PTMs are regulated during this process, we set out to examine the expression of three key tubulin PTMs - acetylation, polyglutamylation and detyrosination - as well as total tubulin levels across neuronal differentiation stages. During differentiation, neuron-specific βIII-tubulin levels and all three PTMs increased following the transition from iPSCs to neurons; however, tubulin acetylation and polyglutamylation showed a markedly steeper increase relative to tubulin detyrosination, or total α-and βIII tubulin (**Figure 1C-D**). This suggests that the enrichment of tubulin acetylation and polyglutamylation are not a consequence of higher tubulin abundance but instead an active regulatory process to support functional and morphological remodeling during differentiation.

**Figure 1.**
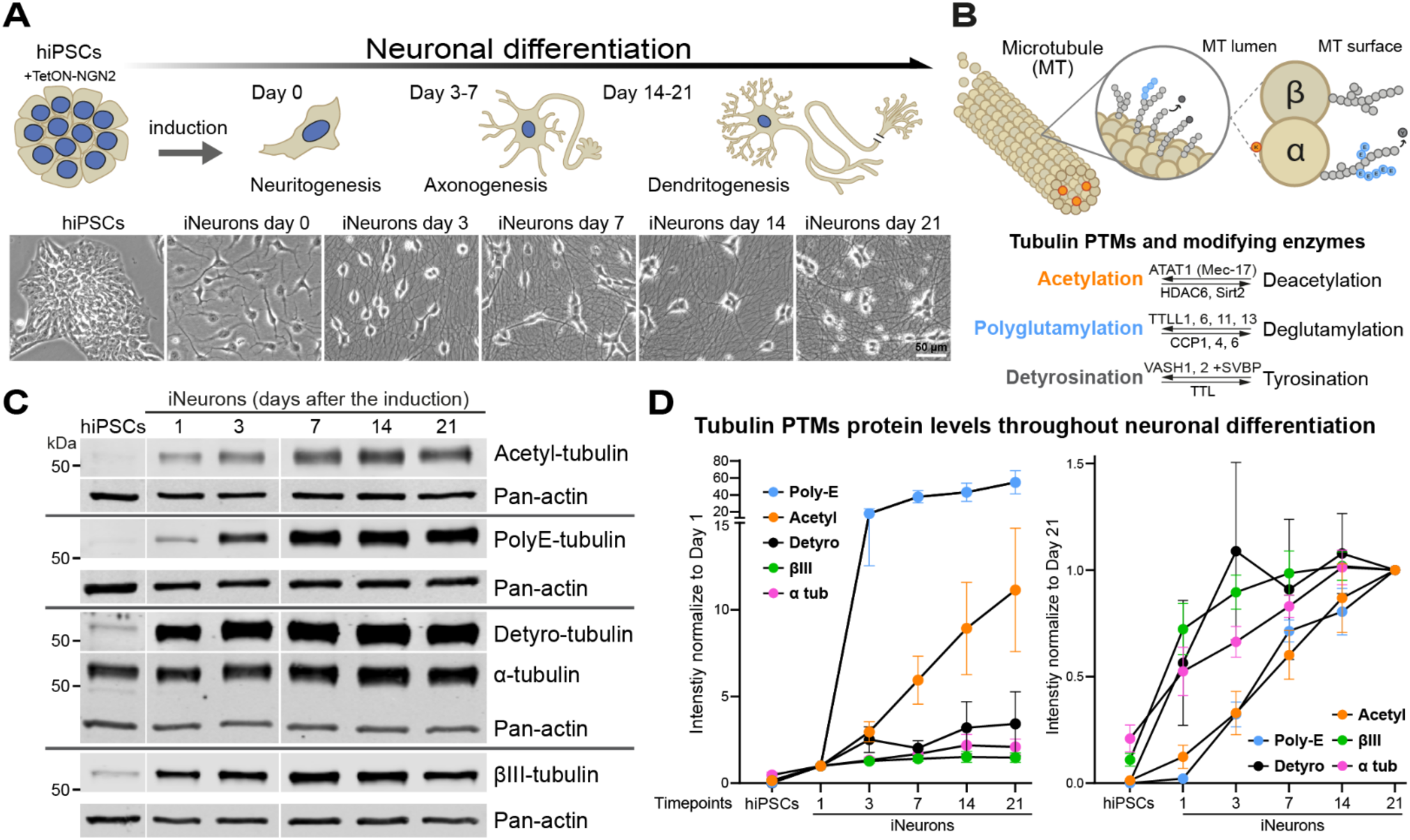
Tubulin PTMs increase in hiPSC-derived cortical neurons during differentiation. (A) Schematic with corresponding brightfield images illustrating the NGN2-induced neuronal differentiation process across distinct morphogenesis stages. Human TetON-hNGN2-KOLF2.1J iPSCs were differentiated into cortical neurons (iNeurons) driven by NGN2 induction. (B) Overview of the three tubulin post-translational modifications (PTMs) examined in this study, modification sites on tubulin, and the known corresponding regulatory enzymes. (C) Western blots of cultured iPSC and iNeuron lysates collected at indicated days of neuronal differentiation, showing the expression patterns of acetylated (Acetyl), polyglutamylated (Poly-E), or detyrosinated (Detyro) tubulin, as well as βlll tubulin, α tubulin, and actin. (D) Quantification of tubulin and tubulin PTM protein levels in iPSCs and iNeurons. The protein levels were first normalized to actin and then normalized to iNeuron day1 (left) or day21 (right) levels. n (culture)=3 per condition. Error bars denote SEM.

### Tubulin PTMs have distinct subcellular distributions during neuronal differentiation

Given the substantial increase in tubulin acetylation and polyglutamylation levels during neuronal differentiation, we next determined where within the cell body these modifications are enriched. To assess their subcellular organization, we methanol fixed iPSCs and iNeurons at different stages of differentiation, stained for each tubulin PTM along with α-tubulin, and performed Airyscan imaging (**Figure 2A**). We qualitatively observed striking differences in PTM distribution between iPSCs, day 0 and day 7 neurons. Acetylated tubulin had a punctate pattern in iPSCs, became filamentous but was restricted to a small subset of microtubules in day 0 iNeurons, and then reached a broader distribution in soma and neurites by day 7 (**Figure 2A**). Polyglutamylated tubulin was not visible in iPSCs, was restricted to the microtubule organizing center in day 0 iNeurons and then took on a broader distribution pattern by day 7 (**Figure 2A**). Tubulin detyrosination displayed a distinct temporal pattern, shifting from a filament-associated distribution at early stages (before day 3) to a punctate distribution along microtubule filaments in more mature neurons (day 7 and 14). Because this partial coating pattern made it difficult to interpret its relationship with microtubule filament organization, we did not include tubulin detyrosination in the following analysis (**Figure S1A**). For quantitative analysis, we measured the intensity ratio of each tubulin PTM relative to total α-tubulin within the soma, divided into three radial bins from perinuclear to periphery. Tubulin acetylation was predominantly enriched in perinuclear and central regions, whereas tubulin polyglutamylation showed a greater enrichment in the peripheral bin (**Figure 2B**). Over differentiation, the perinuclear enrichment of tubulin acetylation gradually decreased with a modest outward distribution into central regions. In contrast, tubulin polyglutamylation showed a progressive increase in the peripheral region (**Figure 2C**), indicating the preparation for high enrichment of polyglutamylated MTs in proximal for neurite outgrowth and branching^48–50^. Together, these results suggest that distinct tubulin PTMs are spatially segregated within neurons, suggesting specialized roles in organizing local MT networks and function in neurons during differentiation.

**Figure 2.**
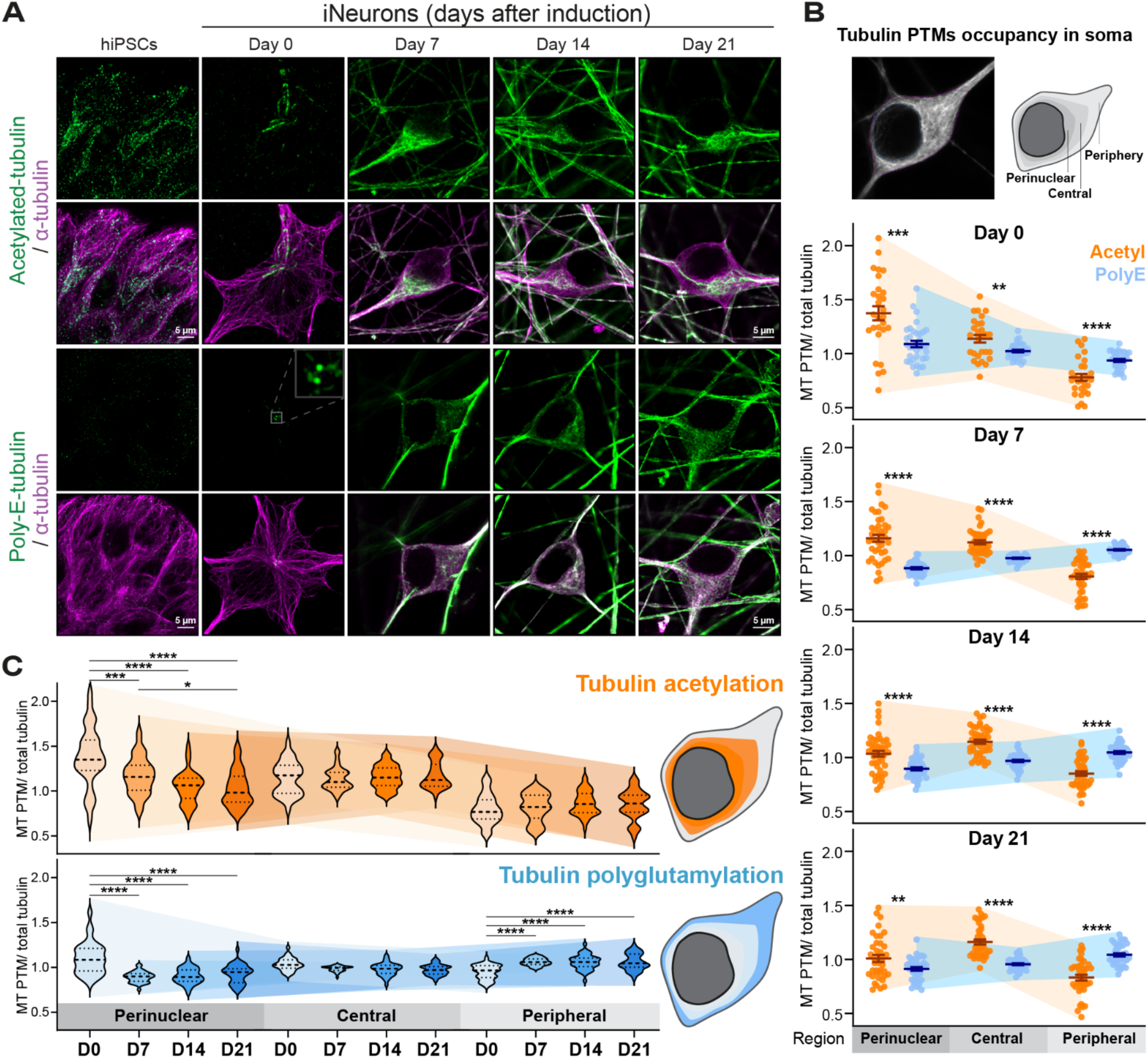
Tubulin PTMs have distinct subcellular distributions during differentiation. (A) Airyscan Z-stack confocal images with maximum intensity projections of methanol fixed iPSCs and iNeurons at day 0, 7, 14, and 21 post induction. Cells were immunolabeled for acetylated or polyglutamylated tubulin (green) and α-tubulin (magenta). (B) Schematic of tubulin PTM occupancy analysis: Z-stack images were Sum intensity projected. Regions of iNeurons were segmented by identifying the soma and nucleus, then radially dividing the soma into three bins, perinuclear, central, and peripheral (Top). The fluorescence intensity of tubulin acetylation or polyglutamylation normalized to total α tubulin was quantified within each bin. Mean intensity ratios per bin are plotted. Two-way ANOVA (Tukey’s multiple comparison post hoc test). **, p<0.01; ***, p<0.001; ****, p<0.0001 are plotted between Acetyl and PolyE groups. n= 28-46 cells per group, pooled from three independent biological replicates. Data are shown as mean ± SEM. (C) Data from (B) were replotted for each tubulin PTM to compare distribution changes across differentiation time points, with comparisons shown as mean ± SEM with one-way ANOVA (Tukey’s multiple comparison post hoc test). *, p<0.05; **, p<0.01; ***, p<0.001; ****, p<0.0001 for tubulin acetylation (Top) and polyglutamylation (Bottom).

### Tubulin acetylation-dependent organelle remodeling occurs predominantly in the soma

Substantive evidence has revealed that multiple membrane-bound organelles are actively remodeled along with metabolic reprogramming as part of the neuronal differentiation process^17,19,21,24^. Our recent work found that remodeling inter-organelle contacts is a hallmark of neuronal differentiation^18^, supporting the transition from pluripotent stem cells to polarized neurons. AnkyrinG staining validated the developmental progression of our iNeuron cultures, indicating that neurons have established polarity with a clearly defined axon on day 7 (**Figure S1B**). The primary enzyme responsible for tubulin K40 acetylation, α-tubulin acetyltransferase 1 (ATAT1)^51^, shows high neuronal enrichment according to human RNA-seq datasets^52,53^ (**Figure S1C**). Consistent with this finding, we observed marked enrichment in tubulin acetylation levels (**Figure 1C and D**) together with upregulation of ATAT1 starting at day 7 and remaining elevated throughout maturation, based on previously published transcriptomics and proteomics^54^ (**Figure S1D**) and Western blotting data (**Figure S1E**). In addition, the specific somatic enrichment of tubulin acetylation suggests a potential regulatory role in coordinating local organelle remodeling during neuronal differentiation. However, how tubulin acetylation governs organelle rearrangements to facilitate their transition from stem cells to neurons remains to be fully delineated. To address this, we set out to reduce tubulin acetylation levels by depleting ATAT1 using an shRNA-based lentiviral approach. Human α-tubulin is encoded by at least nine isotypes^55^ and the acetylatable lysine-40 (K40) residue is conserved among most isoforms except TUBA8. To achieve a broad and isotype-independent reduction in tubulin acetylation, we selectively perturbed ATAT1. Unlike Histone Deacetylase 6 (HDAC6), which deacetylates multiple substrates beyond tubulin, ATAT1 exhibits high specificity for K40 on α-tubulin^56^, providing mechanistically clean intervention for manipulating MT acetylation. When ATAT1 knockdown was initiated in day 3 iNeurons with collection on day 7, the depletion was insufficient. In contrast, shRNA transduction right after NGN2 induction (day 0) followed by collection on day 7 yielded the highest knockdown efficiency in both tubulin acetylation and ATAT1 (**Figure S2A-B**). Even later transduction on day 7 followed by collection on day 14 had least effective suppression, reflecting the slower turnover^57^ of the pre-existing pool of acetylated tubulin (**Figure S2A**). Thus, we selected shRNA transduction on day 0 iNeurons for subsequent experiments.

To investigate how tubulin acetylation mediates organelle networks, we applied a quantitative multispectral microscopy platform to comprehensively capture organelle organization under conditions of tubulin acetylation perturbation. Our multispectral imaging approach overcomes the channel limitations of standard fluorescence imaging and avoids the low throughput, labor intensive, and harsh fixation required for electron microscopy while preserving the speed and sensitivity of conventional confocal microscopy^58^. Multispectral microscopy using a confocal microscope equipped with a spectral detector allows simultaneous imaging 6-8 organelles in live cell^41,43,59^ and we adapted this tool to iNeuron cultures. Furthermore, utilizing the computational analysis pipeline “infer-subc” established in our lab^18,43^, we systematically analyzed 3D organization among eight organelles. Human iPSCs were induced to differentiate into iNeurons and then transduced with either a non-targeting shRNA (shCtrl) or targeting ATAT1 (shATAT1). On the day prior to imaging, iNeurons were transfected with constructs encoding fluorescent organelle markers specific to endoplasmic reticulum (ER), peroxisomes (PO), Golgi (GL), mitochondria (MC), and lysosomes (LS). On day 7, cultured iNeurons were labeled with plasma membrane (PM) and lipid droplet (LD) dyes. Then organelle 3D architecture was captured using multispectral confocal microscopy. Single timepoint 3D Z-stacks were acquired to reconstruct organelle morphology, their inter-organelle communications and spatial distribution (**Figure 3A**). Spectral images were then processed by linear unmixing and deconvolved, followed by instance segmentation of organelles on each intensity channel separately (**Figure S2C-D**). Cell masks were generated from the plasma membrane channel and then subdivided into soma and neurite masks. Nuclear masks were defined by NGN2-BFP2 signal expression. Different mask types were applied to segmented organelle datasets to enable compartment-specific quantification using infer-subc (**Figure 3B** and **Figure S2C-D**). As organelles can exhibit distinct roles, functions, and morphologies between the soma and neuronal processes^17,21,60^, we analyzed soma and neurites separately to capture compartment-specific remodeling. From the analysis pipeline, 5439 metrics per cell were generated and a curated subset was selected to eliminate redundancy, then utilized for statistical comparisons across conditions to screen for differences.

**Figure 3.**
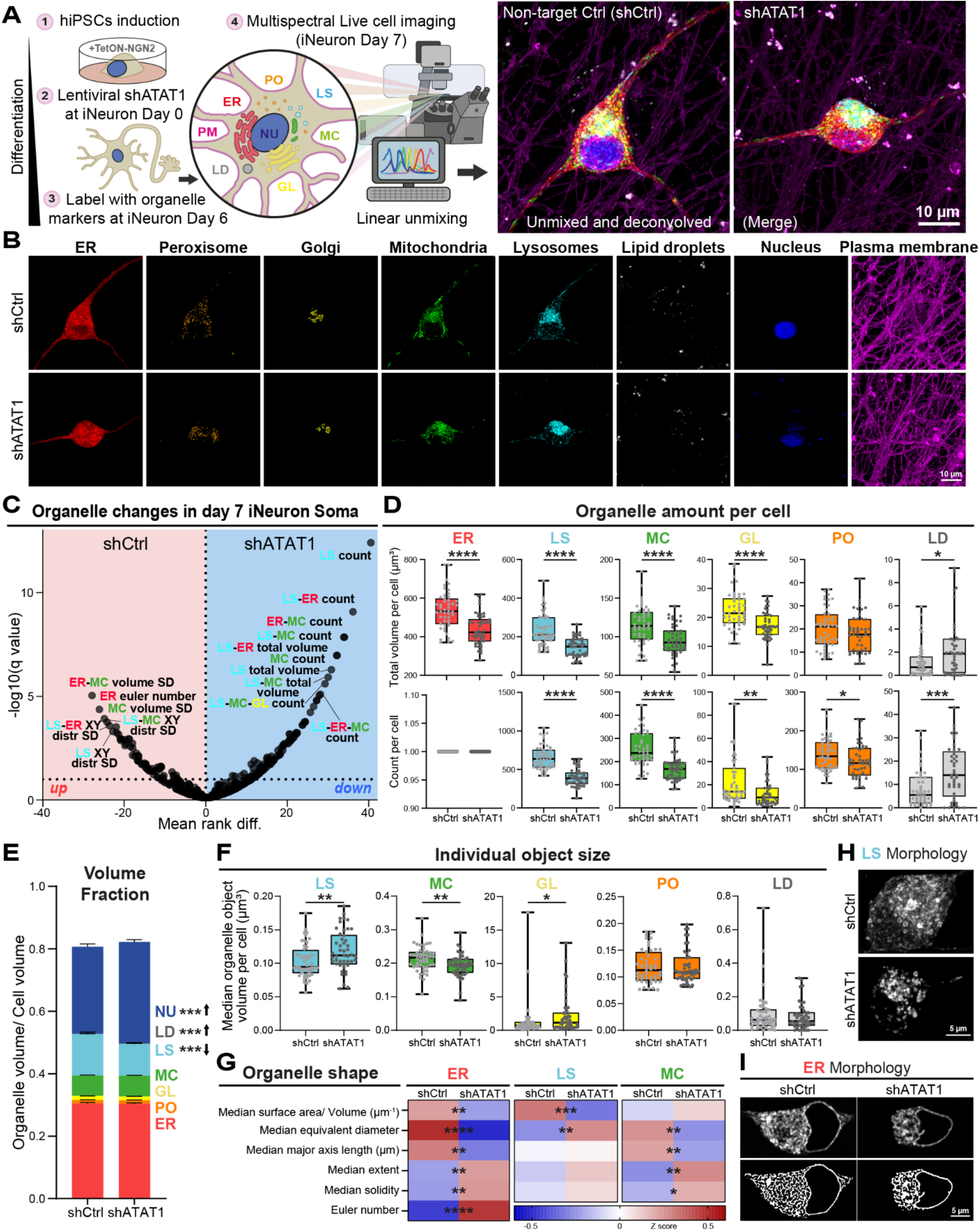
Tubulin acetylation promotes organelle organization during neuronal differentiation. (A) Schematic of multispectral imaging workflow: iNeurons differentiated from hNGN2-hiPSCs were transduced with lentivirus encoding shRNA targeting ATAT1 at day 0. Next, day 6 iNeurons were transfected with 6 different piggyBac plasmids encoding organelle markers in the presence of transposase, and after 24 hours labeled with a lipid droplet fluorescent dye and a plasma membrane stain. All fluorophores were excited simultaneously on a confocal microscope and the emitted light collected using a spectral detector. The overlapping emission spectra was distinguished through linear unmixing, followed by deconvolution using Huygens Essential software. Fluorescent signals were then segmented with an in-house analysis pipeline (Infer-subc) for 3D image analysis. Representative maximum intensity projections of Z-stack multispectral images of shCtrl- and shATAT1-treated day7 iNeurons used for quantitative analysis are shown on the right. (B) Individual channels from processed 3D multispectral images display eight organelles (endoplasmic reticulum (ER), peroxisomes (PO), Golgi (GL), mitochondria (MC), lysosomes (LS), lipid droplets (LD), nucleus (NU), and plasma membrane (PM)) separately. (C) Volcano plot illustrating soma-specific differences in 3D organelle organization (449 metrics) between shCtrl- and shATAT1-treated day7 iNeurons. Statistical significance was determined using a 10% false discovery rate (FDR) threshold. Detailed statistical outcomes and representative metric details are presented in Table S1 and Figure 3 D-G, Figure S3 and S4. SD, standard deviation. distr, distribution. n (cells)=48 shCtrl and 47 shATAT1 from three biological replicates. Asterisks denote q values, * : q < 0.1,** : q < 0.05,*** : q < 0.01,**** : q < 0.001. (D) Box plots display organelle total volume (Top) and number (Bottom) per cell separately in shCtrl-versus shATAT1-treated day 7 iNeurons. Data points represent single cells. The ER was segmented as a single continuous object, resulting in one ER count per cell. (E) Staggered plots show the total individual organelle volume fraction relative to cell volume in each group. Data are presented as mean ± SEM, with asterisks denoting q values from (C). (F) Box plots display individual object median size per cell of five organelles. Asterisks denoting q values from (C). (G) Heatmaps displaying the organelle shape metrics for ER, lysosomes and mitochondria. Each metric is Z-score-normalized with asterisks denoting q values from (C). (H-I) Representative images showing lysosomes (H) and ER (I) in shCtrl- and shATAT1-treated day 7 iNeurons, illustrating morphological differences correlated to quantitative analyses.

We found that depleting tubulin acetylation resulted in distinct compartmental-specific organelle remodeling. A Mann-Whitney U test with multiple comparisons was performed on 449 soma-specific metrics including organelle morphology, organelle interactions and spatial distribution. Applying a false discovery rate (FDR) of 10%, we identified 177 metrics significantly different between shCtrl-and shATAT1-treated day 7 iNeurons (**Figure 3C; Table S1**). A corresponding analysis of neurite regions, including 131 metrics of organelle morphology and interactions, only identified 9 significant differences (**Figure S3A; Table S1**). As human day 7 iNeurons are polarized and highly extended with only limited proximal neurites captured in our imaging field, the soma occupies the majority of total cellular volume in our analysis. Correspondingly, each organelle volume was substantially enriched within the soma compared to neurites (**Figure S3B**). Notably, GL was not detected in neurite regions. Principal component analysis (PCA) based on curated organelle morphology and inter-organelle contacts further demonstrated that soma-derived features can effectively distinguish between shCtrl- and shATAT1-treated groups and provided stronger separation than neurite-specific metrics (**Figure S3C; Table S1**). Together, these findings indicate that depleting tubulin acetylation triggers organelle remodeling predominantly in the soma, suggesting the perinuclear enrichment of tubulin acetylation observed previously plays an important role in organelle organization and positioning in soma during neuronal differentiation. Therefore, our subsequent analyses focused on soma-localized organelle phenotypes, where organelle remodeling was most evident.

### Reducing tubulin acetylation disrupts organelle organization during neuronal differentiation

From the curated 3D organelle analysis, lysosome morphology and lysosome-related organelle interactions were among the most significantly altered features upon tubulin acetylation depletion. We examined overall organelle abundance by quantifying total volume and number per cell and observed that most organelles decreased in both parameters in shATAT1-treated iNeurons (**Figure 3D**). Specifically, the absolute total volume of ER, LS, MC, and GL were lower. In parallel, the number of LS, MC, and PO decreased. Because the ER forms a single continuous network within each cell, only one ER object was segmented per cell. Thus, ER total volume is more representative than its number. When accounting for differences in cell size, only LS showed disproportionate reduction in fraction volume, whereas other organelles scaled proportionally with cell size (**Figure 3E**).

To understand how these abundance changes occur, we next examined the median size of individual organelles and organelle shapes (**Figure 3F, G** and **Figure S3D**). The loss of MC mass arose from combined reduction in number and individual object size. In addition, shape analysis showing decreased median equivalent diameter and major axis length together with increased median extent suggests that MC became fragmented and shorter, losing their elongated tubular morphology. The increase in standard deviation per cell further points to greater variability within the mitochondrial population, consistent with uneven fragmentation (**Figure S3E**). Surprisingly, in contrast to MC, the loss of LS content was mostly due to reduced organelle numbers rather than shrinkage. The remaining LS appeared enlarged, suggesting swelling of remaining LS results from impaired fission and/or enhanced fusion events. The combination shape analysis such as a decreased surface-area-to-volume ratio and increased equivalent diameter revealed that LS became smoother, larger spheres with fewer extensions or tubules. Moreover, the decreased standard deviation of LS volume indicates that the LS population became more uniform, which aligns with visual evidence for the loss of smaller or tubular LS in shATAT1-treated iNeurons (**Figure 3H** and **Figure S3E**). Compared with other organelles, we observed extensive shape remodeling in ER (**Figure 3G** and **Figure S3D**). A decreased surface-to-volume ratio, major axis length and equivalent diameter implies overall shortening and the loss of fine tubular extensions of the ER network. Additionally, increased extent, solidity, and Euler number suggests that the ER network became smoother and more compact. Together, these shape metrics describe ER network in shATAT1-treated iNeurons transitioned from an extended, highly interconnected tubular ER to topologically simplified, less reticulated and sheet-enriched organization (**Figure 3I**). The marked ER shape remodeling is consistent with published studies showing that ER architecture is highly sensitive to acetylated MTs in non-neuronal cells or primary neurons^31,34,61^. In contrast to most other organelles, LDs increased in both total volume, number and fraction volume, but were unchanged in individual size, indicating *de novo* LD formation rather than enlargement of the existing objects in shATAT1-treated iNeurons (**Figure 3D-F**).

Organelle position is closely linked to MT cytoskeleton and is critical for their function^34^. Next, we assessed whether these morphological changes were accompanied by subcellular distribution alterations. We analyzed the spatial distribution of organelles in the lateral (XY) and axial (Z) dimensions within soma. The XY distribution describes the spread from the nucleus towards the soma boundary segmented into five concentric regions, while Z distribution is from the bottom to the top of the soma subdivided into 10 equal bins. Results showed that LS, MC, and GL were distributed closer to the nucleus in shATAT1-treated iNeurons. In addition, the elevated variability of LS and MC lateral distribution suggests an impairment of acetylated MT-dependent transport, leading to a more random organelle positioning (**Figure S3F-G**). On the other hand, LDs exhibited a more outward to periphery distribution, likely reflecting enhanced biogenesis at certain ER subdomains. For axial distribution, we observed most organelles shifted toward the bottom of the shATAT1-treated soma, with a more uniform vertical positioning indicating that perinuclear acetylated MT may guide organelles to extend toward upper soma (**Figure S3F-G**).

Beyond the subcellular phenotypes, we also identified that depleting tubulin acetylation affected overall neuron morphology. We observed a shrinkage of both cell and soma size, but neurite number was not altered (**Figure S3H**). In summary, acetylated tubulin-deficient day 7 iNeurons display distinct organelle morphological and distribution changes, as well as changes in overall neuron morphology. Generally, we found reduced abundance of LS that were enlarged, rounder, and centrally clustered.

The ER became less reticulated and MC more disorganized and fragmented.

### Tubulin acetylation governs organelle interactions during neuronal differentiation

Contact sites between membrane-bound organelles are increasingly recognized as critical determinants of neuronal architecture and homeostasis^62^. Supporting this idea, recent work has shown that organelle interaction patterns can distinguish neurons from other cell types, such as astrocytes^43^. The remodeling of organelle morphology and positioning can strongly influence their engagement with each other. However, no studies have yet systematically characterized how tubulin acetylation regulates these inter-organelle interactions during neuronal differentiation. Through quantitative 3D inter-organelle profiling, we identified distinct patterns of organelle interaction remodeling following ATAT1 knockdown. We quantified organelle interactions by measuring the overlapping regions between organelle objects, referred to as organelle interaction sites or “contacts”. Although this approach does not directly discriminate between membrane contacts (10-80 nm) and close proximity (<410 nm) processes like co-trafficking, our prior work demonstrated that interaction-site measurements closely serve as a reliable proxy for *bona fide* membrane contacts, as detected with dimerization-dependent contact biosensors^63^. The analysis revealed that upon tubulin acetylation depletion, changes in interaction abundance were most pronounced among pairwise organelle interactions, with fewer three-way interactions and very limited higher-order (>3-way) interactions affected (**Table S1**). Among all analyzed organelles, ER-, LS- and MC-related interactions were the most abundant pairwise interactions (**Figure 4A**). Depleting tubulin acetylation, we found an overall reduction in interaction volume and number per cell among these organelles, whereas PO-related interactions were less affected. For example, 4/5 ER-organelle, 4/5 LS-organelle, and 3/5 MC-organelle contact numbers were reduced in ATAT1-depleted cells (**Figure 4A**). In contrast, the least abundant contacts, those involving LDs, were increased in shATAT1-treated iNeurons, likely due to increased LD biogenesis.

**Figure 4.**
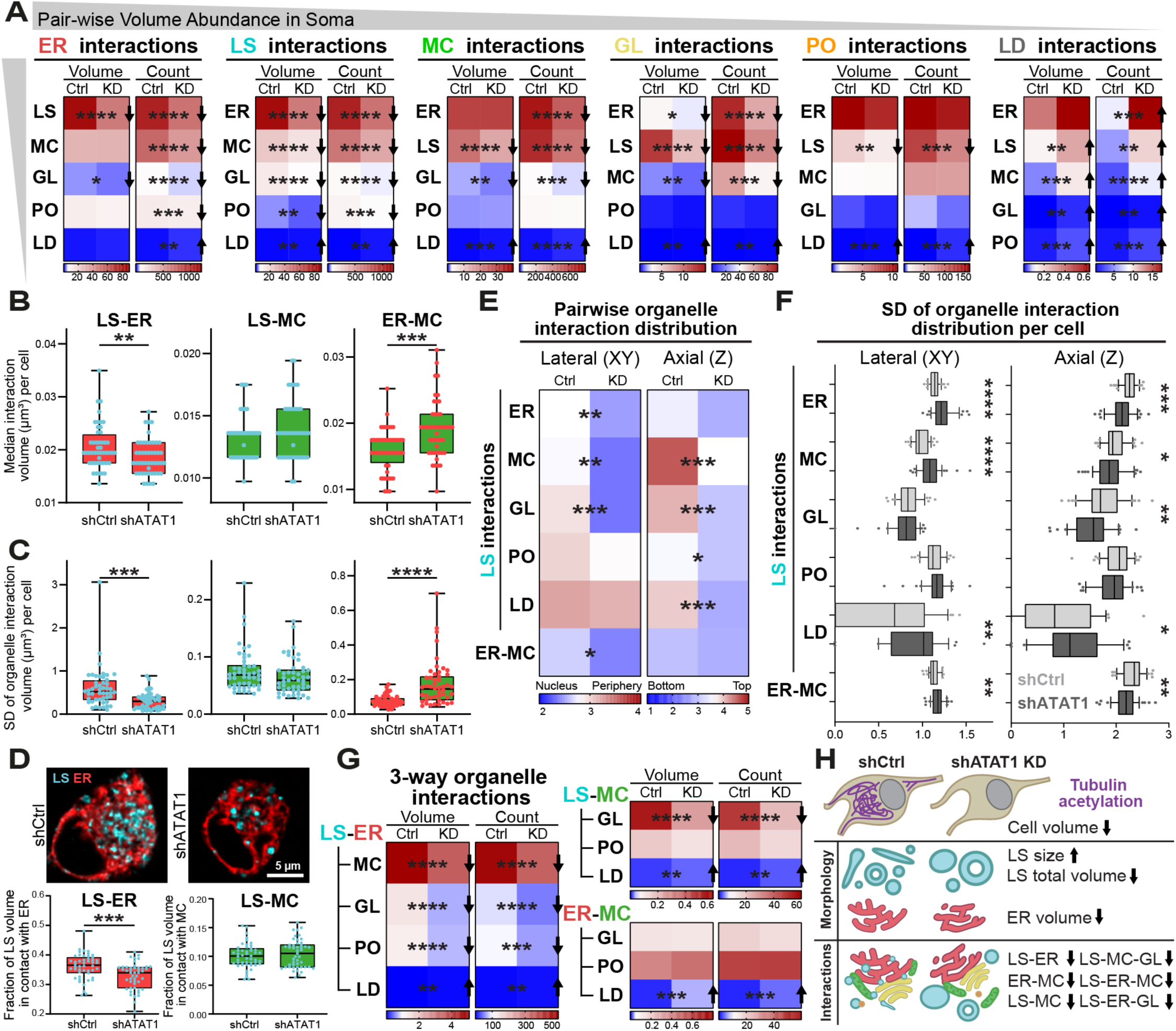
Tubulin acetylation governs organelle interactions during neuronal differentiation. (A) Heatmaps of the total volume (μm^3^) and number of pairwise soma-specific organelle interactions per cell in shCtrl-versus shATAT1-treated day 7 iNeurons, ordered by organelle volume abundance. Asterisks indicate q-values from Figure 3C. n (cells)=48 shCtrl and 47 shATAT1 from three biological replicates. Arrow direction indicates whether the value increases or decreases in shATAT1 relative to shCtrl. (B-C) Median individual interaction volumes per cell for the three most abundant organelle pairs (lysosome-ER, lysosome-mitochondria and ER-mitochondria) are shown in (B) and the corresponding standard deviation (SD) of organelle interaction volumes per cell is shown in (C). Asterisks indicate q-values from Figure 3C. n (cells)=48 shCtrl and 47 shATAT1 from three biological replicates. (D) Representative images of lysosomes (cyan) and ER (red). The fraction of lysosomes contacting ER (bottom left) or mitochondria (bottom right) relative to total lysosome number per cell was quantified for the proportion of lysosomes engaged in ER or mitochondria interactions. Statistical significance was determined using an unpaired t-test with ***, p<0.001. (E) Heatmaps summarize the mode lateral or axial distribution metrics for lysosome-related and ER-mitochondria interactions from the soma-specific 3D multispectral organelle analysis. (F) Corresponding boxplots show the variability (SD, standard deviation) of pairwise organelle interaction distributions along the lateral and axial axes. (G) Heatmaps of the volume (μm^3^) and number of soma-specific three-way organelle interactions among lysosome, ER and mitochondria per cell in shCtrl-versus shATAT1-treated day 7 iNeurons. Asterisks indicate q-values from Figure 3C and n (cells)=48 shCtrl and 47 shATAT1 from three biological replicates in (E-G). (H) Schematic summarizing the key organelle morphology and interaction changes identified in shCtrl-versus shATAT1-treated day 7 iNeurons through multispectral 3D organelle analysis, with interaction order reflecting the ranking of interaction changes from the volcano plot Figure 3C.

Among all interaction changes, the number of LS-ER interactions exhibited the greatest reduction in count as well as volume, followed by the number of ER-MC and LS-MC in shATAT1-treated iNeurons (**Figure 3C**). Notably, ER-MC contacts have been shown to occur on acetylated MTs in non-neuronal cells^31^, which is consistent with our finding that depleting tubulin acetylation results in disrupted ER-MC contacts. Our results suggest a similar acetylated tubulin-mediated mechanism operates during neuronal differentiation. ER-MC interactions exhibited a significant reduction in number accompanied by greater variability and increased interaction size, while total interaction volume/cell remained unchanged, suggesting impaired interaction initiation but compensation by enlarging or stabilizing the remaining interactions (**Figure 4A-C**). Meanwhile, LS-MC interactions decreased in both total volume and number, but were unchanged in interaction size or variability, indicating the reduction was primarily due to fewer encounters and unchanged interaction structure (**Figure 4A-C**). LS-ER interactions in shATAT1-treated iNeurons displayed reductions across number, total and individual interaction volume, along with decreased variability of interaction volume/cell (**Figure 4A-C**). Most importantly, when considering the proportion of LS engaged in interactions, we identified a significant decrease in the fraction of LS volume in contact with ER, while the fraction of LS volume in contact with MC was unchanged (**Figure 4D**). This indicates broad and uniform LS-ER interaction disruption, rather than decreased LS-ER interactions being merely a consequence of fewer organelles being present. Similarly, line-scan intensity profiles show greater distance between ER and LS in shATAT1-treated iNeurons, suggesting weaker coupling (**Figure S4A**). The collective reduction of LS-ER interaction metrics suggest that LS-ER interactions are the most sensitive to acetylated tubulin depletion. A similar reduction was also detected in proximal neurites (**Figure S3A**).

Following the global reduction in organelle interactions, we next examined their spatial distribution in both lateral (XY) and axial (Z) dimensions of the soma. Most LS-related contacts as well as ER-MC and GL-related interactions became more enriched towards nucleus in shATAT1-treated iNeurons. In the axial dimension, most pairwise organelle contacts shift downward. However, LS-ER and ER-MC relatively maintained their vertical position (**Figure 4E** and **Figure S4B**). Additionally, we found several LS-related organelle contacts were laterally dispersed yet vertically confined toward the basal region in shATAT1-treated iNeurons (**Figure 4F**). Minimal changes in distribution variability were observed among other organelle pairs (**Figure 4C**).

The most frequent 3-way interactions involved interaction between the ER, LS and MC, which were also the most abundant pairs in the 2-way interactions. Among 3-way interactions, we identified the amount of multiple LS-ER-organelle interactions (LS-ER-MC, LS-ER-GL, and LS-ER-PO) showed a significant reduction following ATAT1 knockdown, as well as observing a reduction in LS-MC-GL and LS-GL-PO interactions (**Figure 4G** and **Figure S4D**). In summary, beyond changes in organelle abundance and morphology, depleting tubulin acetylation led to extensive remodeling of organelle interactions and their spatial distribution, with the most pronounced effects on LS-ER contacts as well as higher-order interactions involving LS-ER. The key changes upon ATAT1 knockdown are summarized (**Figure 4H**).

### Lysosome-ER contacts preferentially associated with acetylated tubulin

From the quantitative 3D organelle screening, we identified lysosome-ER (LS-ER) interactions as the most affected by tubulin acetylation depletion. While the confocal-based multispectral analysis provided systematic and comprehensive quantification, to visualize with higher spatial resolution whether LS-ER interactions are positioned on acetylated MTs, we next employed a 3-colored STED super-resolution imaging. Cells were fixed and endogenously labeled for ER (calnexin), total α-tubulin, acetylated tubulin and lysosomes (LAMP1) in shCtrl-versus shATAT1-treated iNeurons. In addition to the day 7 timepoint, we also examined day 14 iNeurons to assess the effects of prolonged tubulin acetylation depletion (**Figure 5A** and **Figure S5A**). Effective acetylated MT depletion was observed at both timepoints (**Figure S5B**). Additionally, sustained depletion of ATAT1 for 14 days did not result in widespread cell death or severe degeneration, though the neuronal network appeared less elaborate with fewer processes (**Figure 5A**). Consistent with multispectral results, we observed enlarged lysosomes in ATAT1 knockdown cells, which became even more pronounced after 14 days of ATAT1 depletion (**Figure 5B**). LS-ER contacts displayed a decreasing trend in day 7 shATAT1-treated iNeurons, aligning with our earlier findings, and the decrease became more evident after prolonged depletion (**Figure S5C**).

**Figure 5.**
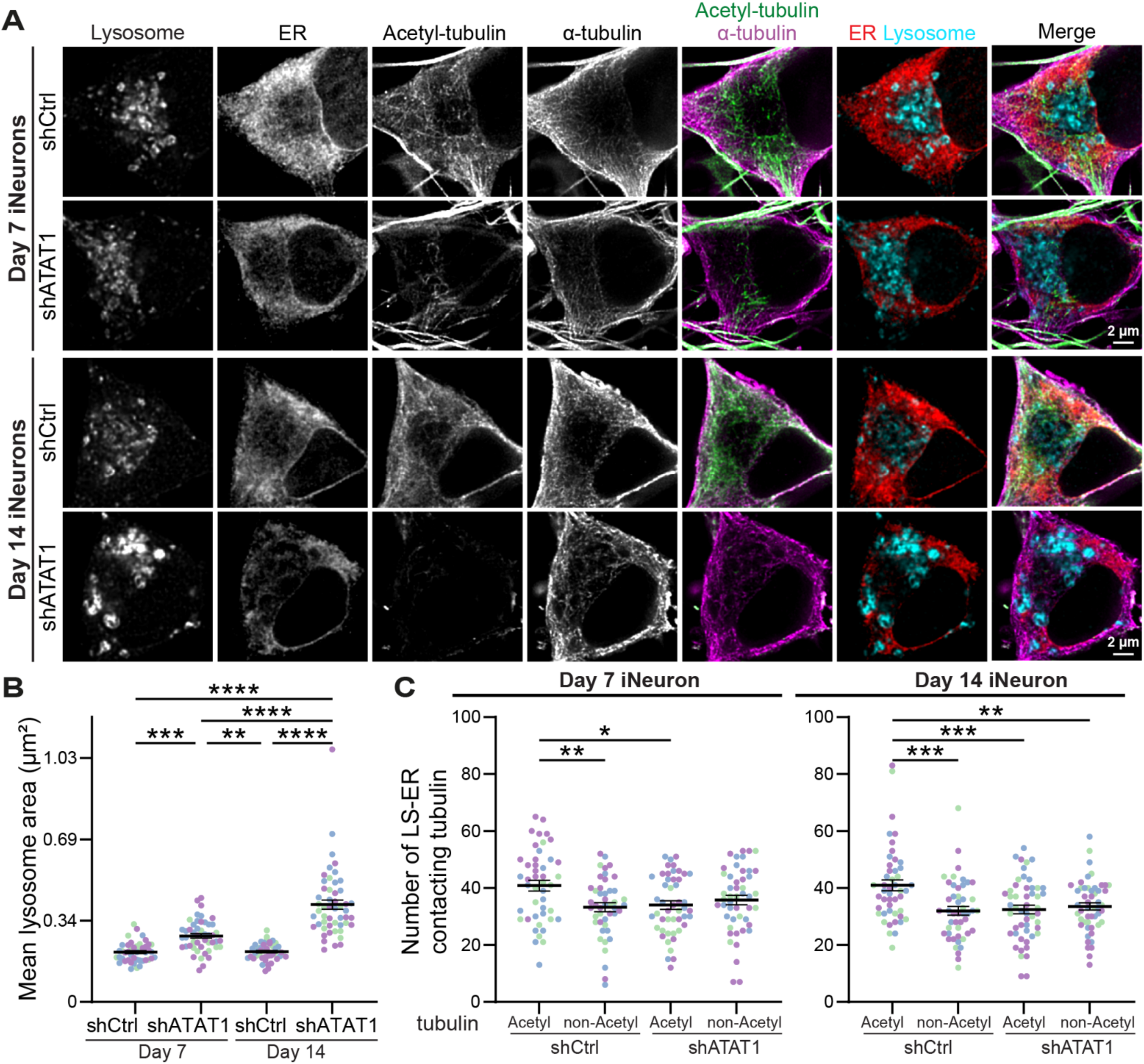
Super-resolution analysis reveals that lysosome-ER contacts preferentially associate with acetylated tubulin. (A) Three-color stimulated emission depletion (STED) imaging of shCtrl-versus shATAT1-treated iNeurons fixed on day 7 or day 14, illustrating subcellular interaction of lysosomes with ER in nanoscale. ER, acetylated and total tubulin were visualized in STED mode using antibodies against calnexin, acetyl K40 tubulin and α tubulin, respectively. Lysosomes were imaged in confocal mode and labeled with a LAMP1 antibody. (B-C) Individual fluorescence channels from (A) were segmented using CellProfiler to create object masks for quantification. Non-acetylated tubulin masks were generated by subtracting acetylated tubulin masks. Lysosome mean size was quantified from the segmented images and compared between each condition (B). The number of lysosome-ER contact sites overlapping with acetylated tubulin or non-acetylated tubulin regions were identified from segmented masks quantified in (C). Data represent mean ± SEM with one-way ANOVA (Tukey’s multiple comparison post hoc test). n (cells)=45-51 per group from three biological replicates, with data points color-coded by replicate. *, p<0.05; **, p<0.01; ***, p<0.001; ****, p<0.0001.

We next examined whether LS-ER contacts preferentially associated with acetylated MTs in soma. In shCtrl-treated iNeurons, both the number and total area of LS-ER contacts overlapped significantly with acetylated MTs compared to non-acetylated MTs. However, in neurons depleted of tubulin acetylation, LS-ER contacts associated with acetylated tubulin were reduced, while LS-ER contacts associated with non-acetylated tubulin were unaffected (**Figure 5C** and **Figure S5D-F**). This indicates that the reduction in observed LS-ER contact sites was due to loss of enrichment of these contacts on acetylated MTs in ATAT1-depleted cells. This preference was consistently observed on both day 7 and day 14 iNeurons. Together, these results indicate that tubulin acetylation provides a specialized MT framework to spatially facilitate the formation and organization of LS-ER interactions during neuronal differentiation.

### Tubulin acetylation promotes lysosomal fission in developing neurons

LS-ER contacts have been shown to function as key interfaces to regulate lysosomal morphology and dynamics. Lysosomes exhibit distinct dynamics when engaged in ER tubule contacts along MTs compared with free transport, suggesting the idea that MT-based lysosome dynamics vary with ER association^64^. Studies in non-neuronal cells showed that upon induced lysosomal damage, lysosomes form tubular extensions and ER tubules are often present before the fission of extended lysosome tubules^65^. Additionally, shifting ER morphology to more ER sheets decreases lysosomal fission^65^. In neurons, ER tubules control lysosome size, mediate their fission, promote kinesin-1 loading onto late endosomes/ lysosomes within contact region^66^ and facilitate lysosome entry into axon^67^. The reduced LS-ER contact abundance we observed in acetylated tubulin-deficient iNeurons could thus cause a disruption of lysosomal dynamics, particularly affecting lysosomal tubulation.

We hypothesized that altered lysosomal tubulation events could explain the enlarged lysosomal morphology observed in shATAT1-treated iNeurons, as impaired tubule fission would decrease the formation of smaller lysosomes. To investigate this possibility, we next examined lysosomal dynamics through live-cell imaging and quantified tubulation events. We classified the lysosomal tubules into three categories, retractile, stable and fission. Over three minutes of time-lapse images, lysosomes that extended and then retracted without forming new vesicles were counted as retractile tubules. Stable tubules remained long-lasting and steady in morphology. Lysosomal tubules that had new vesicles released from the parent lysosomes were grouped as fission events (**Figure 6A** and **Supplementary Video 1**). Though the tubulation events were not significantly different between shCtrl- and shATAT1-treated iNeurons, tubulated frequency was lower as neurons matured (**Figure S6A**). Upon tubulation analysis, we found that more than half of lysosomes were fission type, followed by retractile tubules in shCtrl-treated iNeurons. However, this balance was flipped in shATAT-treated iNeurons, with a greater fraction of retractile than fission events observed in neurons after 7 days or 14 days of ATAT1 knockdown. This result suggested that tubulin acetylation is required for lysosomal fission throughout different neuronal differentiation processes (**Figure 6B** and **Figure S6B**). The amount and fraction of lysosomal tubulation events associated with mature lysosomes were quantified using cathepsin B activity sensor (Magic Red) (**Figure S6C-D**). The shift from fission to retractile lysosomal tubules together with reduced LS-ER interactions indicates an impairment of lysosomal scission mediated by ER tubules. The failure of lysosomal fission can prevent small lysosomal regeneration and deplete the total lysosome pool, leading to progressive lysosome enlargement.

**Figure 6.**
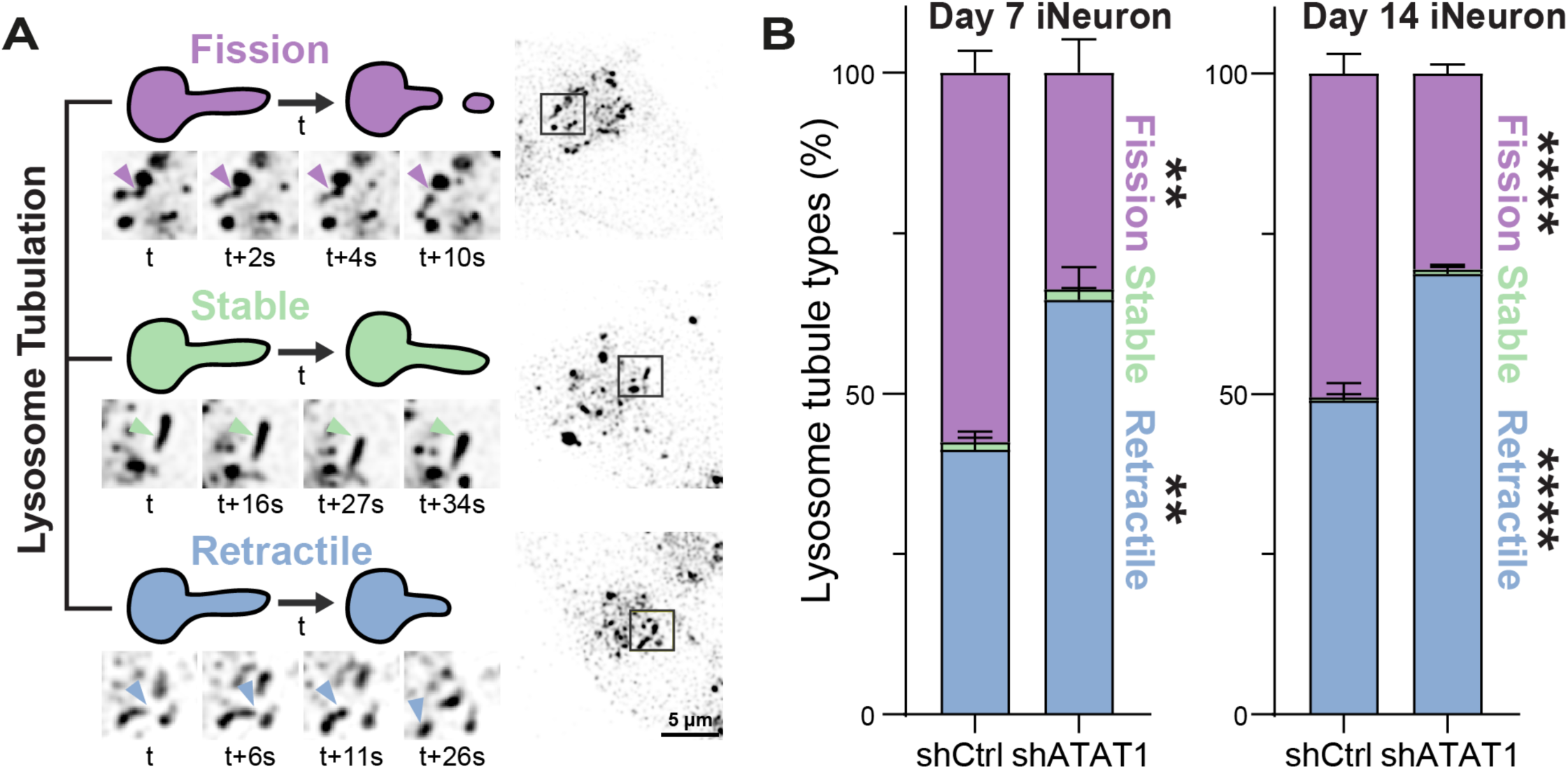
Tubulin acetylation promotes lysosomal fission. (A) Time-lapse confocal imaging showing dynamics of lysosomes labeled with Lysotracker-Green and imaged on day 7 or day 14 for 3 minutes (1 frame/s) in shCtrl- versus shATAT1-treated iNeurons. Lysosome tubulation events were quantified and categorized as fission (purple), stable (green), or retractile (blue). Representative images show fission, stable, or retractile lysosome tubules originating from the same lysosome in day 7 shCtrl-treated iNeurons. Arrowheads indicate the site of the event. The time-lapse images on the left correspond to the gray boxed region on the right. (B) Staggered graph showing the percentage frequency of different lysosome tubule types quantified within the soma. Statistical significance between shCtrl- versus shATAT1-treated groups for each tubule type is shown with data representing mean ± SEM with two-way ANOVA (Šidák’s multiple comparison post hoc test). n (cells)=74 day 7 shCtrl, 64 day7 shATAT1, 138 day14 shCtrl and 101 day14 shATAT1 from three biological replicates. *, p<0.05; **, p<0.01; ***, p<0.001; ****, p<0.0001.

### Loss of tubulin acetylation increases lysosomal acidity and promotes autolysosome accumulation

Lysosomes are known for nutrient sensing, maintaining iron homeostasis, autophagy, and intracellular signaling. Their acidic lumen is critical for activating hydrolases for macromolecule degradation. Enlarged or swollen lysosomes, depending on the pathway perturbed, are often accompanied by alterations in lysosomal acidity^68–70^. Therefore, we next measured lysosomal acidity using a dual-color fluorescence assay sensitive to luminal pH changes to determine whether the lysosomal defects are coupled with changes in lysosomal function. To quantify changes in lysosomal acidity, we measured both the ratio of acidic lysosomal signal intensity to total lysosomal mass, and the total acidic area within the soma of shCtrl-versus shATAT1-treated iNeurons. Surprisingly, we found a significantly increased ratio in ATAT1 knockdown iNeurons compared to the shCtrl cells, suggesting more acidified lysosomes. Consistently, the total area occupied by acidic lysosomes was higher in shATAT1-treated iNeurons among both timepoints (**Figure 7A**). To be noted, more mature neurons (day 14) displayed a greater acidity intensity ratio and total area than younger neurons (day 7), whereas individual acidic lysosome area was altered only in day 14 shATAT1-treated iNeurons (**Figure S7A**). Additionally, when applying an alternative acidity indicator, we observed similar trends that older neurons contained more acidic compartments, and loss of tubulin acetylation further increased the amount of acidic compartments. Moreover, extended depletion of tubulin acetylation (21 days) produced comparable effects to those seen at day 14 (**Figure S7B**). The presence of active luminal lysosomal protease Cathepsin B and Cathepsin D indicates that lysosomes are still capable of undergoing maturation even with defective morphology and dynamics in acetylated tubulin-deficient iNeurons (**Figure S7C**).

**Figure 7.**
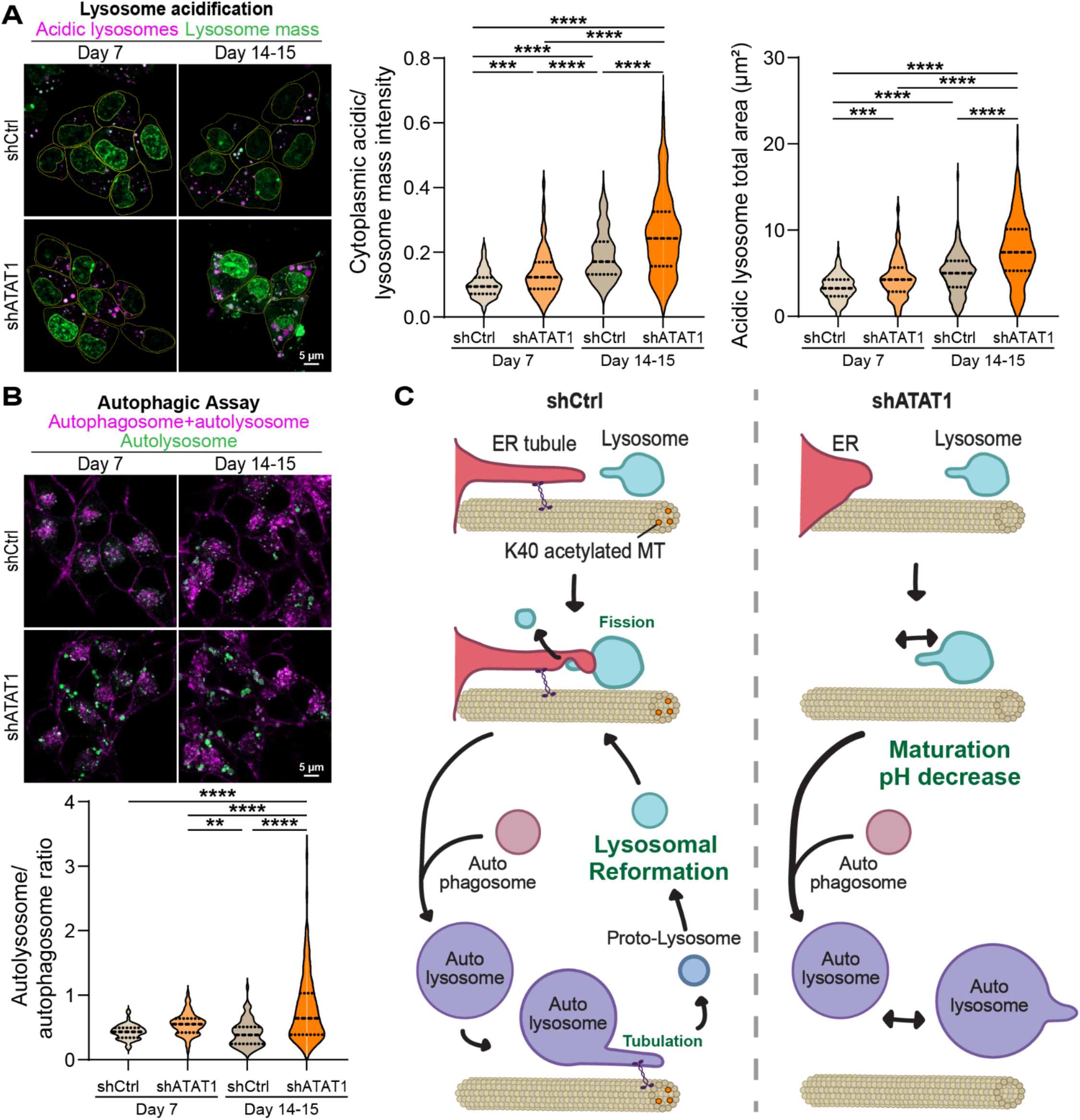
Tubulin acetylation alters lysosome function. (A) Representative confocal images of shCtrl-versus shATAT1-treated iNeurons stained with the lysosomal acidic pH detection kit and imaged on day 7 or day 14-15 (left). The fluorescence intensity ratio of total (green) to acidic (magenta) lysosomes was quantified within the soma (yellow lines) for each condition (middle). Acidic lysosome signals were segmented and analyzed to determine the total acidic lysosome area per soma (right). n (cells)=193 day 7 shCtrl, 170 day7 shATAT1, 185 day 14-15 shCtrl and 154 day14-15 shATAT1 from three biological replicates. Statistical significance was determined using one-way ANOVA (Tukey’s multiple comparison post hoc test) *, p<0.05; **, p<0.01; ***, p<0.001; ****, p<0.0001. (B) Representative confocal images of shCtrl-versus shATAT1-treated iNeurons stained with autophagic assay kit and imaged on day 7 or day 14-15 (top). The fluorescence intensity ratio of total autophagosome + autolysosomes (green) to autolysosomes (magenta) was quantified within the soma for each condition (bottom). n (cells)=62 day 7 shCtrl, 92 day7 shATAT1, 164 day 14-15 shCtrl and 143 day14-15 shATAT1 from three biological replicates. Statistical significance was determined using one-way ANOVA (Tukey’s multiple comparison post hoc test) *, p<0.05; **, p<0.01; ***, p<0.001; ****, p<0.0001. (C) Schematic model summarizing the proposed mechanism of tubulin acetylation-dependent regulation of organelle arrangement and lysosome function during neuronal differentiation.

The coexistence of enlarged, more acidic lysosomes and defects in fission suggest a failure to complete the lysosomal reformation process required to replenish the lysosomal pool. We hypothesized such defects would result in an exhausted autophagic-lysosomal system and accumulation of autolysosomes. Thus, next we examined whether autophagic progression is altered upon tubulin acetylation depletion. We quantified the intensity ratio of autophagosomes and autolysosomes using a live cell autophagic assay. We found ATAT1 knockdown iNeurons, particularly day 14, displayed greater accumulation of autolysosomes relative to total autophagic structures (**Figure 7B**). Overall, these results suggest that 7 days of depletion of tubulin acetylation causes early defects in lysosomal recycling and turnover. By day 14, this phenotype became more pronounced, with lysosomes showing further enlargement, increased acidification, and accumulation of autolysosomes. These patterns indicate a transition from initial lysosome recycling defects to a failure in autophagic lysosome reformation, leading to the persistence of accumulated terminal degradative compartments.

## Discussion

In this study, we define the role of α-tubulin K40 acetylation in organelle remodeling for neuronal differentiation. We first discovered that tubulin acetylation was substantially enriched in differentiating neurons compared with total tubulin. We also found its catalytic enzyme ATAT1 increases in abundance during this process, suggesting active establishment of the modification as neurons mature. We further identified a distinct perinuclear population of acetylated tubulin that accumulated over the course of neuronal differentiation. The position of this perinuclear acetylated MT pool suggests that it functions as a strategic cytoskeletal hub capable of coordinating the organization of diverse organelles. Despite previous characterization of tubulin acetylation as a regulator of axonal transport within neurite regions^34,35^, its role in remodeling organelle organization such as by shaping organelle architecture and organelle interactions within neuronal soma was under explored. Moreover, most previous studies connecting MT PTMs to organelle interactions were performed in non-neuronal cells^31,42,71,72^. Here we identified that depletion of ATAT1 reduced tubulin acetylation levels, especially the soma population, and resulted in widespread multi-organelle remodeling across ER, peroxisome, Golgi, mitochondria, lysosomes, and lipid droplets in day 7 iNeurons, as revealed by our multispectral 3D organelle analysis pipeline. We also demonstrated that tubulin acetylation drives compartment-specific organelle remodeling, influencing organelles in both the soma and neurites. This study provides the first systems-level characterization of organelle organization in live human neurons following manipulation of tubulin acetylation. Our multispectral 3D imaging pipeline quantified 5,439 metrics per cell, capturing organelle morphology, multi-organelle interactions and spatial distribution from Z-stack images. A refined set of soma-specific metrics was sufficient to distinguish control from acetylated MT-depleted iNeurons and uncovered subtle but coordinated organelle changes induced by acetylated MT depletion. Reduction of tubulin acetylation resulted in striking ER remodeling in the soma, with 3D shape metrics revealing a shift from a highly intertwined and branched ER network toward a more sheet-like structures. These changes align with prior evidence that ER morphology is strongly shaped by MTs particularly acetylated MTs^73^ that co-distribute with tubular ER in neurons^34^. Our findings also correlate with known mechanisms by which ER tubules slide along acetylated MTs^31^. Moreover, recent work showing that extracellular cues can induce ER tubulation by activating ATAT1 further supports the idea that acetylated MTs can drive dynamic ER remodeling^61^.

The tubular ER network forms extensive contact sites with other organelles. Thus, the reduction in ER tubules that we observe aligns with the decrease in ER-organelle contacts identified in our analysis. ER tubules mediate lysosome fission in neurons^67^ and during lysosome repair^65^. Loss of ER extension along acetylated MTs is therefore predicted to impair lysosomal fission. In agreement, lysosomes in acetylated tubulin-depleted iNeurons predominantly generated retractile tubules rather than completing fission. Super-resolution STED imaging identified predominant lysosome-ER contacts occurring on acetylated MT enriched regions, supporting a direct spatial relationship between tubulin acetylation and lysosome-ER coupling.

Overall, our findings support a model in which ER tubules extend along acetylated MTs to encounter lysosomes and facilitate their fission. When acetylated MTs are depleted, lysosomes fail to undergo efficient fission and instead continue to mature, acidify, and eventually fuse into autolysosomes. This leads to a reduced number of autolysosomes and lysosomes that are larger, rounder, and more uniform in size. Consistent with the known observation that autolysosomes are typically larger than lysosomes^74^, our data suggest that lysosomes progressively merge into but fail to escape the autolysosomal pool. Normally, autophagic lysosome reformation (ALR) enables autolysosomes to generate new lysosomes through KIF5B-driven extension of autolysosomal tubules, which then undergo scission^75^. Prior work has shown that KIF5B preferentially transports lysosomes along acetylated α-tubulin-enriched perinuclear MTs in HeLa cells^32^, raising the possibility that reduced acetylation limits the availability of these preferred tracks. Although we did not directly evaluate KIF5B function, the buildup of large autolysosomes in acetylated tubulin-depleted iNeurons strongly suggests an impairment in ALR and a block in supplementing the new lysosome pool (**Figure 7C**). Importantly, our results introduce tubulin acetylation as a previously unrecognized regulator of lysosome fission and reformation. We do not exclude the possibility that ER tubules contribute to ALR for lysosome regeneration but exploring this mechanism falls beyond the scope of this study.

In summary, our study highlights the K40 acetylated MT network as a central organizer of organelle remodeling during neuronal differentiation, revealing how neurons utilize cytoskeletal modification to reshape multi-organelle communication. Specifically, tubulin acetylation emerges as an upstream regulator of lysosome homeostasis that acts through changes in lysosome-ER interactions and ALR capacity.

## Supplemental information

**Figure S1.**
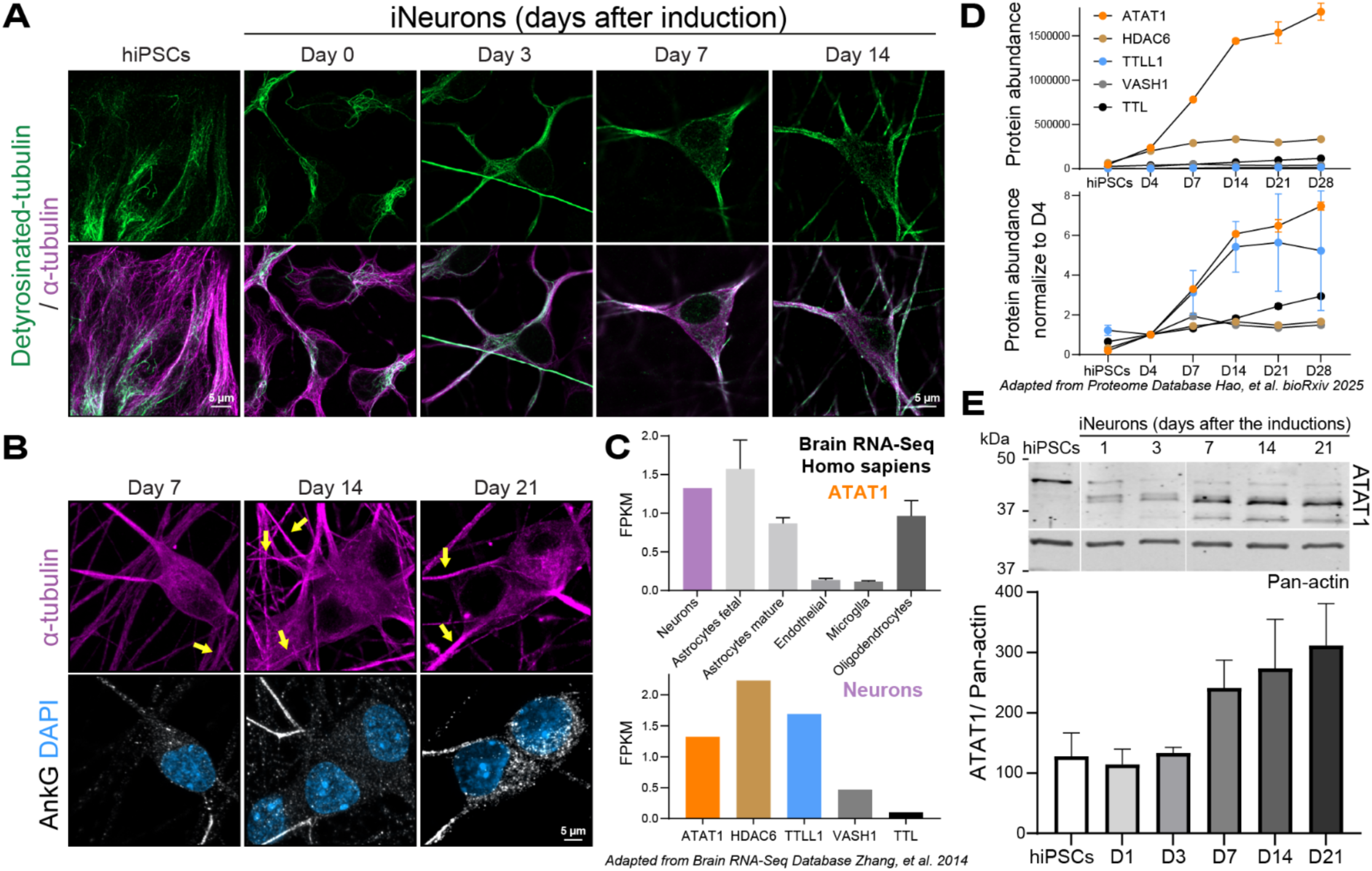
Spatiotemporal regulation of tubulin PTMs and PTM enzymes during neuronal differentiation. (A) Airyscan Z-stack confocal images with maximum intensity projections of iPSCs and iNeurons at day 0, 3, 7 and 14 post induction. Cells were methanol fixed at indicated days and immunolabeled with antibodies against detyrosinated tubulin (green) and α-tubulin (magenta). (B) Representative images of day 7, 14 and 21 iNeurons fixed with methanol and stained for AnkyrinG (AnkG), α-tubulin, and DNA (DAPI) to demonstrate that from day 7 axons (yellow arrows) from iNeurons are specified. (C) RNA-seq results adapted from the Brain RNA-seq Database (Zhang et al., 2014), demonstrating that ATAT1 is expressed in multiple brain cell types, with high levels in human neurons (top). Expression patterns of tubulin PTM-related enzymes in neurons are shown below. FPKM: Fragments Per Kilobase of transcript per Million mapped reads. (D) Protein expression results adapted from the mass spectrometry-based proteomic database (Hao et al., bioRxiv 2025), showing tubulin PTM enzyme enrichment dynamics across iNeuron differentiation. (E) Representative and quantitative western blot analysis of ATAT1 expression pattern in cultured iPSCs and iNeurons collected at indicated days of neuronal differentiation. n (culture)=3 per condition. Error bars denote ±SEM.

**Figure S2.**
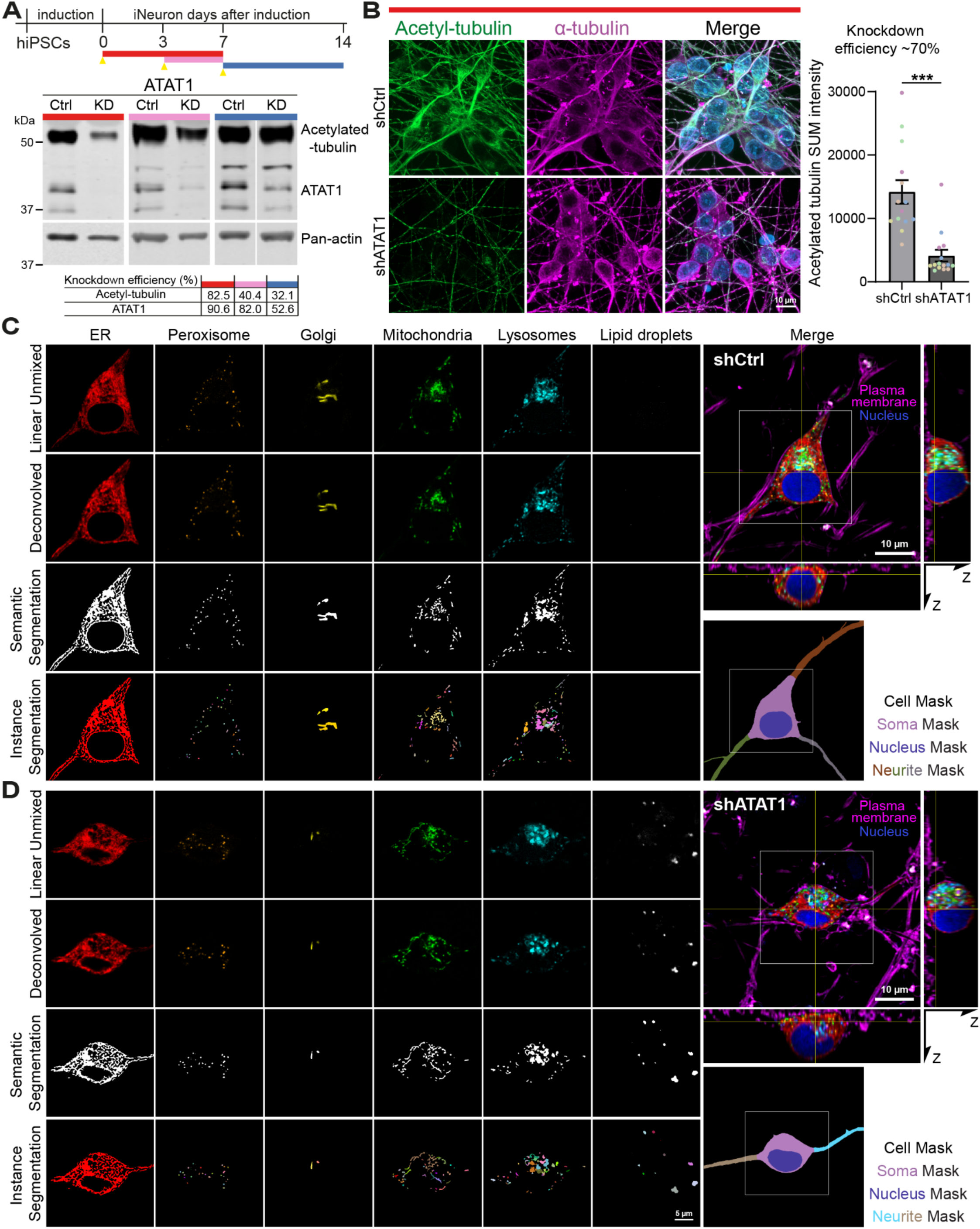
ATAT1 knockdown validation and application of multispectral image pipeline for quantitative organelle analysis in acetylated-tubulin-depleted iNeurons. (A) Schematic diagram of non-target shRNA (shCtrl) versus ATAT1 (shATAT1) lentivirus transduction and cell lysate collection timelines, where yellow arrowheads mark the day of viral transduction and bars denote the duration from transduction to sample collection. Three different timelines were tested, and western blot analysis shows correlating ATAT1 and acetylated tubulin protein levels. Knockdown efficiency was calculated by normalizing the signal intensity of ATAT1 or acetylated tubulin to pan-actin and then comparing the normalized values between knockdown and control groups. (B) Representative immunofluorescence confocal images show shCtrl-versus shATAT1-treated day7 iNeurons fixed and stained for acetylated tubulin (green), α-tubulin (magenta) and DNA (DAPI, blue). Bar graph shows the quantification of tubulin acetylation SUM projected signal intensity from five biological replicates (color-coded), demonstrating decreased tubulin acetylation in shATAT1-compared with shCtrl-treated iNeurons. Upaired two-tailed t test. Error bars represent ±SEM. ***, p<0.001. (C-D) Representative images illustrating the multispectral processing steps: Linear unmixed fluorescence image after spectral separation, deconvolved images using Huygen deconvolution software, segmentation outputs and cell masks using infer-subc pipeline of shCtrl (C) and shATAT1 (D) day7 iNeurons.

**Figure S3.**
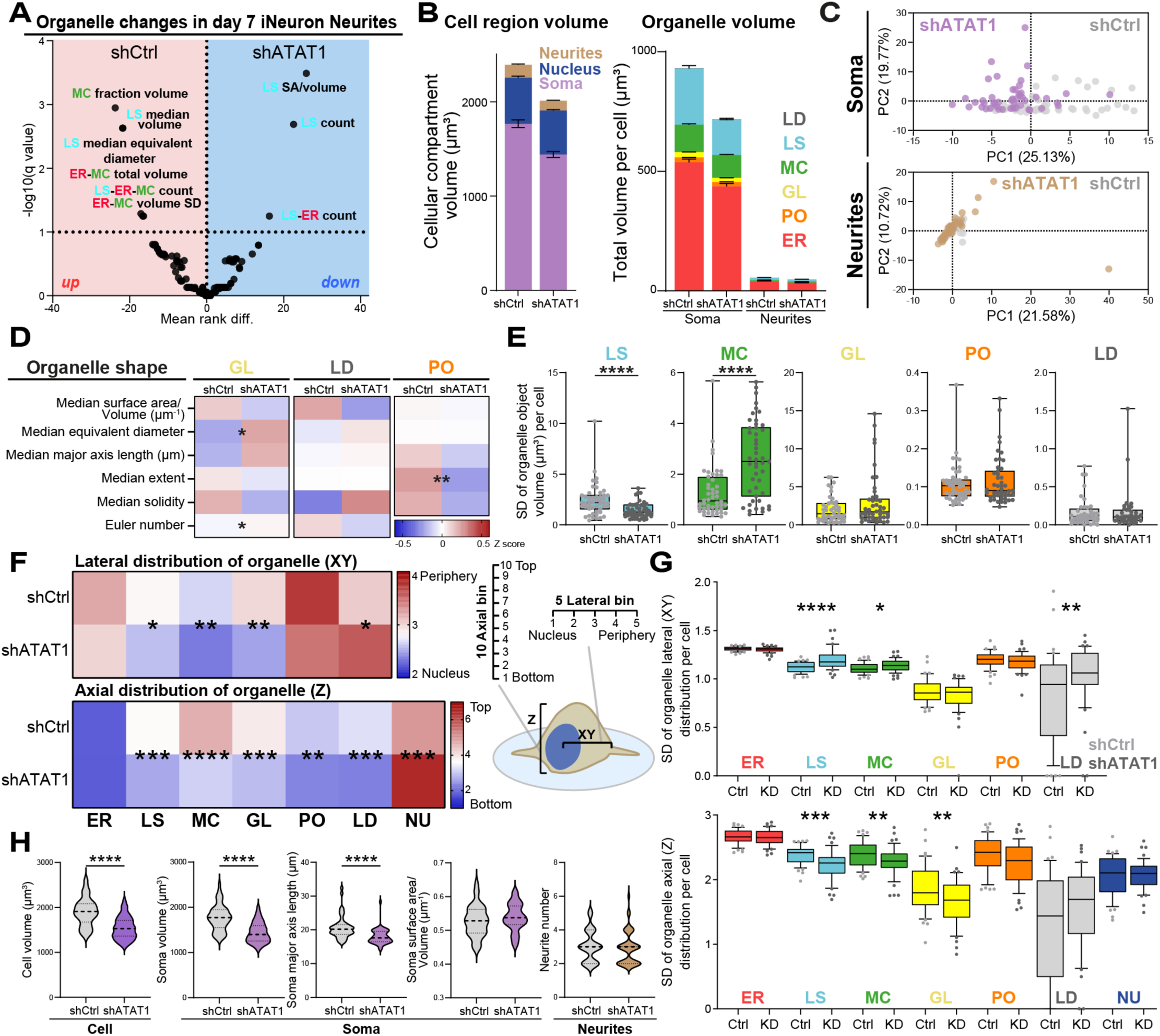
Depletion of tubulin acetylation drives changes in organelle distribution and cell morphology. (A) Volcano plot illustrating neurite-specific differences in 3D organelle organization (131 metrics) comparing shCtrl-versus shATAT1-treated day7 iNeurons. Statistical significance was determined using a 10% false discovery rate (FDR) threshold. SD, standard deviation. SA, surface area. n (cells)=48 shCtrl and 47 shATAT1 from three biological replicates. (B) Staggered plots show cell region volumes for the soma, neurites, and nucleus (left) and each organelle volume within soma and neurites (right). Data are presented as mean ± SEM. (C) Principal component (PC) scores based on PC analysis of 3D organelle morphology and interaction metrics (248 soma-specic metrics; 131 neurite-specific metrics) reveal a clear separation between shCtrl and shATAT1 day 7 iNeuron soma. The percentages on each axis represent variance explained by the PC. Data points represent single cells. (D) Heatmaps displaying the soma-specific organelle shape metrics for Golgi, lipid droplets and peroxisomes. Each metric is Z-score-normalized with asterisks denoting q values from Figure 3C. (E) Boxplots show standard deviation (SD) values of organelle object volumes for each organelle per cell. Asterisks denote q values from Figure 3C. n (cells)=48 shCtrl and 47 shATAT1 from three biological replicates. (F) Schematic of organelle distribution measurements along the XY and Z axes (right). For lateral (XY) distribution, segmented Z-stacks were summed across Z planes and divided into five equal bins with region 1 as the nucleus and regions 2-5 spanning from the nuclear edge to the cell periphery. Object volumes of six organelles within each bin were quantified and the mode values of each organelle were displayed as a heatmap (top left). For axial (Z) distribution, summed projections across XY axes were divided into ten equal bins from bottom to top (1 to 10) to quantify organelle volumes within each Z-section, with a heatmap displaying the mode value of six organelles plus nucleus across Z dimension (bottom left). Asterisks denote q values from Figure 3C. (G) Boxplots of lateral (top) or axial (bottom) distribution standard deviation (SD) values for each organelle. Asterisks denote q values from Figure 3C. n (cells)=48 shCtrl and 47 shATAT1 from three biological replicates. (H) Quantification of cell morphology analysis including cell and soma volumes, soma major axis length, surface-to-volume ratio and neurite number between shCtrl-versus shATAT1-treated day 7 iNeurons; statistical significance was determined using an unpaired t-test with **, p<0.01; ***, p<0.001; ****, p<0.0001.

**Figure S4.**
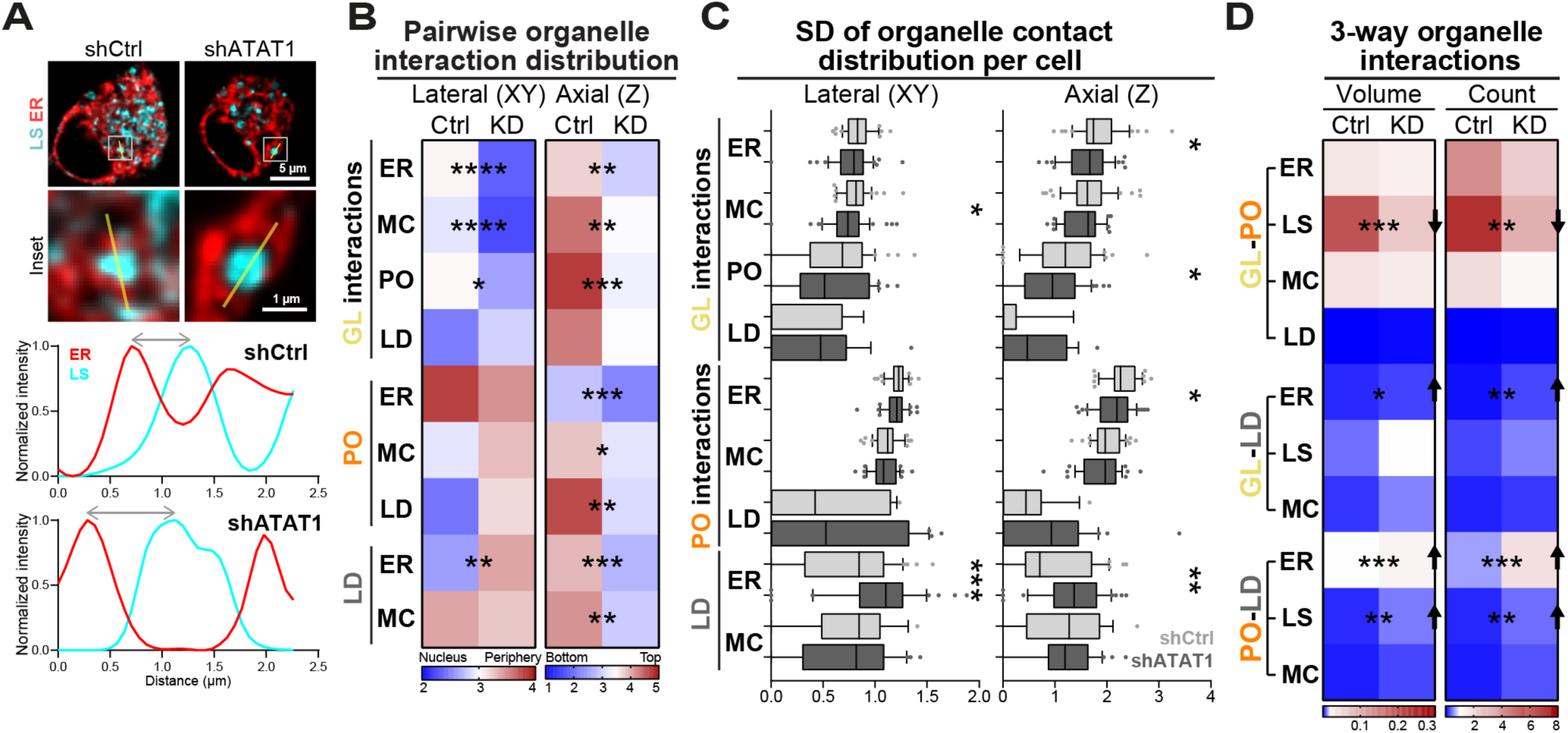
Extended organelle interaction profiling under tubulin acetylation depletion during neuronal differentiation. (A) Line scan profiles showing lysosome-ER distance in shCtrl-versus shATAT1-treated day 7 iNeurons, with normalized intensities measured across the yellow lines. (B-D) Heatmaps in (B) show the mode lateral or axial distribution metrics of soma-specific pairwise interactions among Golgi, peroxisome and lipid droplets per cell in shCtrl- versus shATAT1-treated day 7 iNeurons. The corresponding standard deviation (SD) of interaction volumes per cell is shown in (C). Heatmaps of the number and volume of soma-specific three-way organelle interactions among Golgi, peroxisome and lipid droplets per cell are shown in (D). Asterisks indicate q-values from Figure 3C and n (cells)=48 shCtrl and 47 shATAT1 from three biological replicates in (B-D).

**Figure S5.**
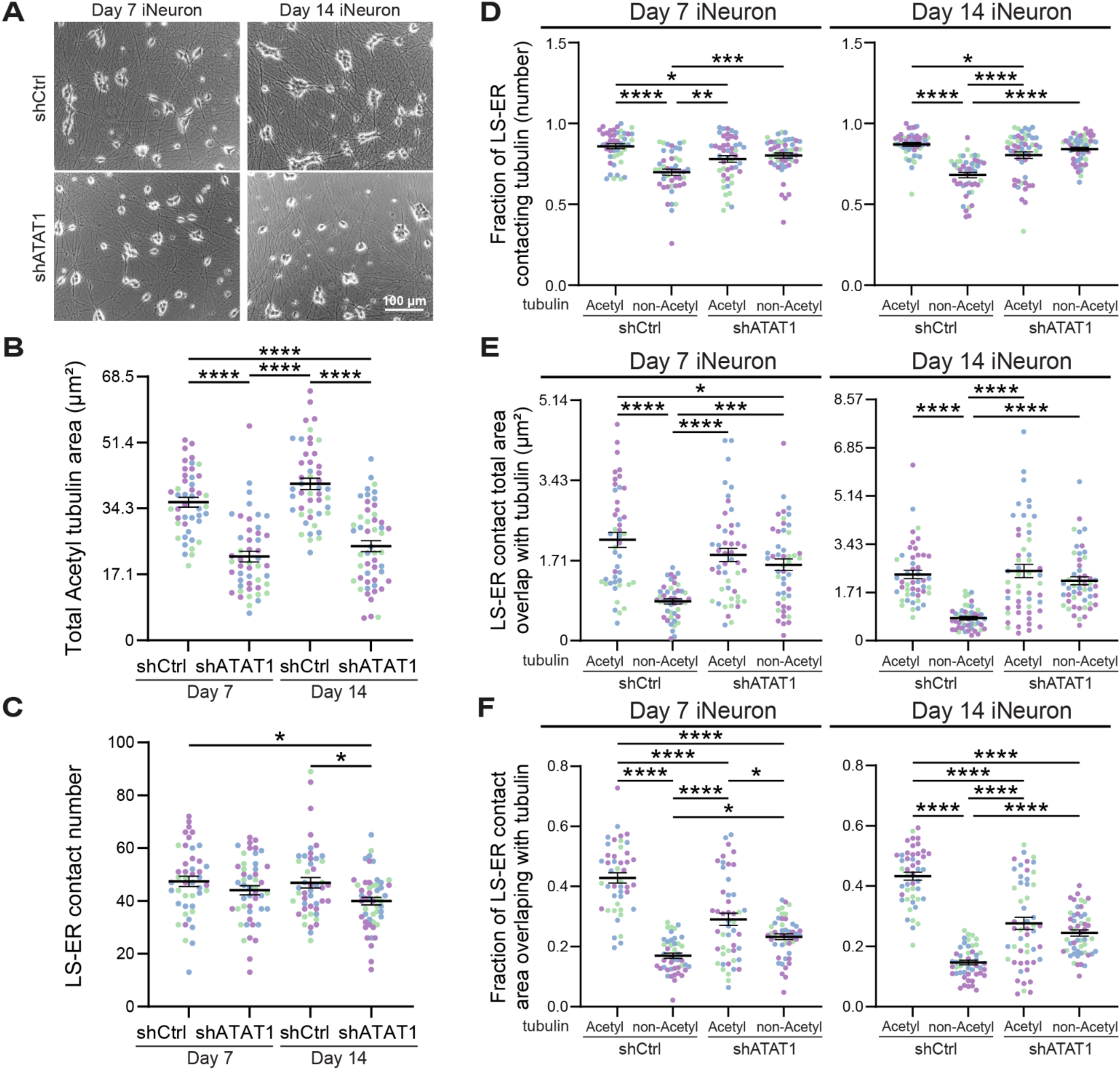
STED analysis reveals lysosome-ER contact area is enriched on acetylated tubulin. (A) Brightfield images of iNeuron cultures following 7 days (left) or 14 days (right) of ATAT1 knockdown. (B) Total acetylated tubulin area per cell, quantified from the segmented images and compared between conditions. (C) Number of lysosome-ER contacts quantified by counting overlaps between segmented organelle masks. (D) The fraction of lysosome-ER contact number colocalized with each tubulin type relative to total lysosome-ER contact number per cell. (E) The area of lysosome-ER contact sites overlapping with acetylated tubulin or non-acetylated tubulin regions identified from segmented masks. (F) The fraction of lysosome-ER contact area colocalized with each tubulin type relative to total lysosome-ER contact area per cell. In (B-F), data represent mean ± SEM with one-way ANOVA (Tukey’s multiple comparison post hoc test). n (cells)=45-51 per group from three biological replicates, with data points color-coded by replicate. *, p<0.05; **, p<0.01; ***, p<0.001; ****, p<0.0001.

**Figure S6.**
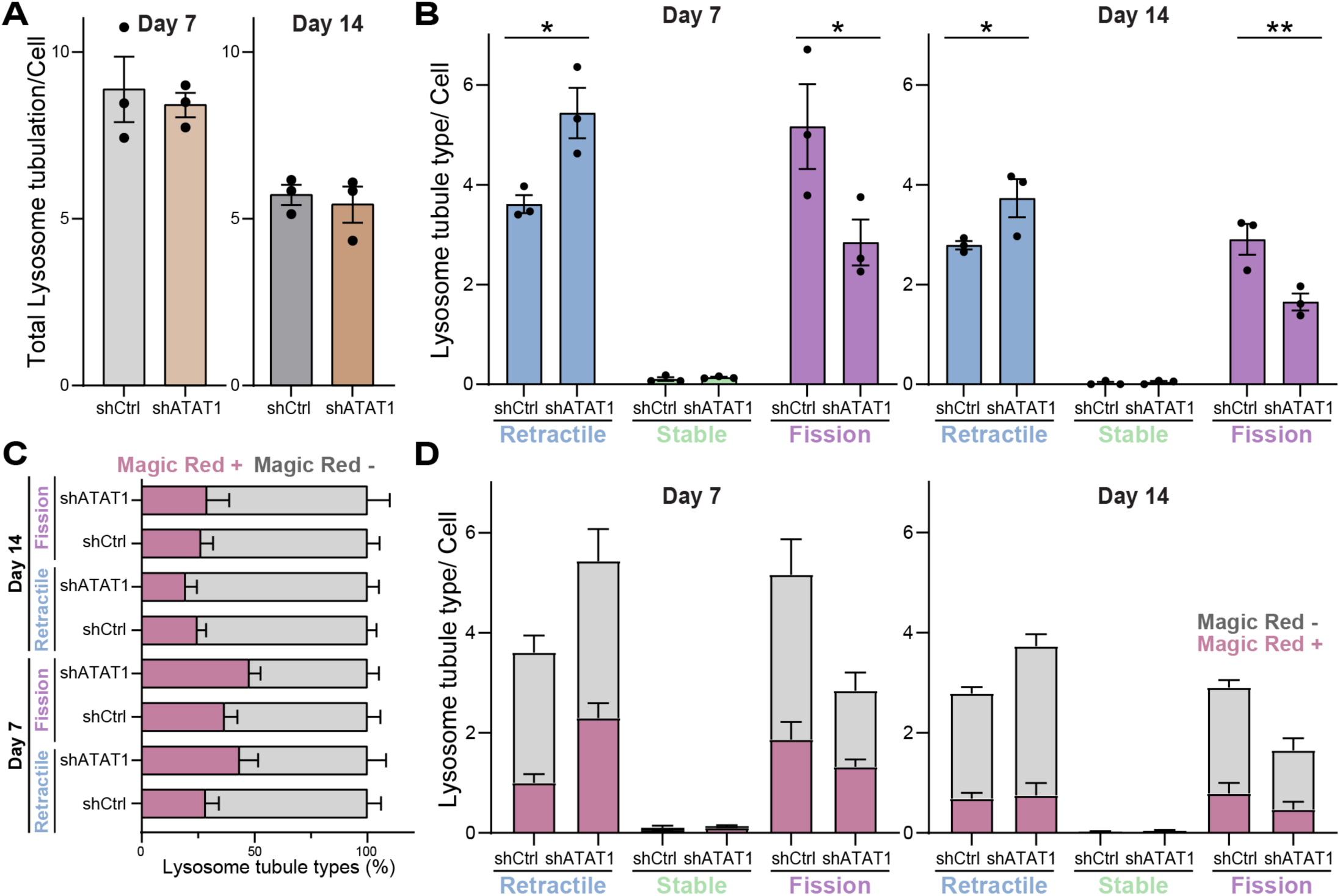
Analysis of lysosomal tubulation events under reduced tubulin acetylation. (A) Same conditions as Figure 6, plotted here as total tubulation event number per cell in shCtrl- versus shATAT1-treated iNeurons. n (replicate)=3. (B) Number of each tubulation event type per cell (fission, stable, retractile). Statistical significance between shCtrl-versus shATAT1-treated groups for each tubule type is shown with data representing mean ± SEM with two-way ANOVA (Šidák’s multiple comparison post hoc test). n (replicate)=3. *, p<0.05; **, p<0.01; ***, p<0.001; ****, p<0.0001. (C) Staggered graph showing the percentage of Magic Red positive or negative lysosome tubule types quantified within the soma. (D) Numbers of each tubulation event type per cell (fission, stable, retractile), further divided into Magic Red positive or negative pools.

**Figure S7.**
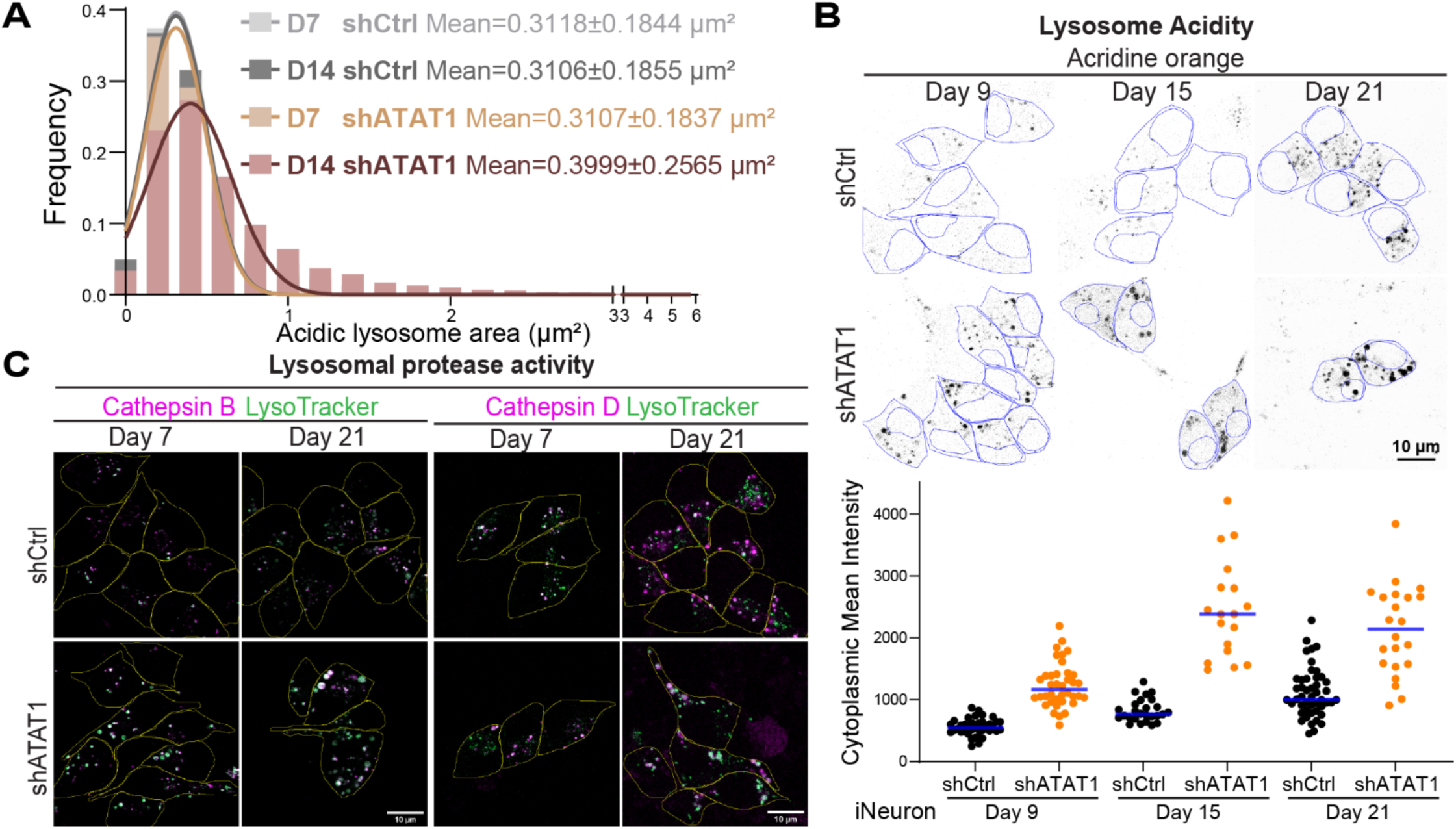
Depleting tubulin acetylation does not disrupt lysosome maturation. (A) Frequency histograms of individual segmented acidic lysosome area from day 7 shCtrl (n=193 cells), day7 shATAT1 (n=170 cells), day 14-15 shCtrl (n=185 cells) and day14-15 shATAT1 (n=154 cells) from three biological replicates. (B) Inverted confocal images of shCtrl-versus shATAT1-treated iNeurons stained with acridine orange to label acidic compartments and imaged on day 9, day 15 or day 21. Quantification of acridine oranges means intensity per soma (blue) shows higher acidic compartment signal in shATAT1-treated neurons, with the effect becoming more evident following prolonged knockdown. (C) Lysosomal protease activity, assessed using Magic Red (cathepsin B) and SiR-Lyso (cathepsin D), is observed in enlarged lysosomes in both day 7 and day 21 iNeurons, indicating that enlarged lysosomes remain enzymatically active during prolonged ATAT1 knockdown.

**Supplementary Video 1:** The video shows a cropped region of lysosomal tubulation dynamics corresponding to the boxed area in Figure 6A.

## Methods

### Culture of human iPSC derived neurons

Human KOLF2.1J wildtype (wt) induced pluripotent stem cells (hiPSCs) were obtained from Bill Skarnes (The Jackson Laboratory) and were used to generate a pBac-TetON-hNGN2 iPSCs stable cell line as previously described^18,44^. Briefly, KOLF2.1J wt hiPSCs were transfected with 500 ng piggyBac plasmid encoding a bipartite TetON-hNGN2 cassette (plasmids gifted from Michael Ward; NINDS). Following transfection, cells were selected for stable genomic integration using puromycin (0.7 µg/ml). Cells were maintained and cultured as adherent monolayers at 37°C in a humidified incubator with 5% CO_2_ under feeder-free conditions in StemFlex medium (A3349401; Gibco) on vitronectin (VTN-N)-coated 6-well plates (VTN-N Recombinant Human Protein, Truncated, A14700; Gibco). Plates were coated with vitronectin by diluting 10 µL vitronectin in 1 mL of 1x Dulbecco’s Phosphate-Buffered Saline (DPBS, 14190144; Gibco) per well and incubating for 2 hrs at 37°C. The hiPSCs were media changed every 1-2 days and were passaged upon reaching ∼80% confluency using ReLeSR (#100-0484; STEMCELL Technologies) according to manufacturer’s instructions in the presence of 0.5x ROCK inhibitor (RevitaCell Supplement, A2644501; Gibco).

### Differentiation into iNeurons

An established protocol was adapted to differentiate pBac-TetON-hNGN2 iPSCs into induced neurons (iNeurons)^38,44^, with modifications^18^. Cultured hNGN2-iPSCs were dissociated into single cells using Accutase (A6964-100ML; Sigma-Aldrich) and cells (1.0-1.2×10^6^ per well of 6-well plates) were seeded in induction medium (IM) consisting of KnockOut™ DMEM/F-12 (12660012; Gibco) supplemented with 1x N-2 (17502048; Gibco), 1x MEM Non-Essential Amino Acids Solution (MEM-NEAA, 11140050; Gibco), 1x GlutaMAX (35050061; Gibco), 1x ROCK inhibitor (RevitaCell Supplement, A2644501; Gibco) and 0.5 μg/ml Doxycycline hyclate (D9891; Sigma-Aldrich). Twenty-four hrs after plating, replaced media into IM (1x N-2, 1x MEM-NEAA, 1x GlutaMAX) with 1 μg/ml Doxycycline hyclate. After 48 hrs, 4×10^4^ induced cells were replated using Accutase onto each well of 8-well Nunc Lab-Tek II chambers with 0.1mg/mL poly-L-ornithine hydrobromide (Sigma; #P3655) in Borate buffer (10 mM Boric acid (Sigma B6768), 2.5 mM Sodium tetraborate (Sigma 221732), 7.5 mM Sodium chloride (Sigma S7653), 0.1 M Sodium hydroxide (Sigma 71463)) and 10 μg/mL Laminin (Gibco™ Laminin Mouse Protein, Natural, 1mg; 23-017-015) coated in IM (1x N-2, 1x MEM-NEAA, 1x GlutaMAX) with 0.5 μg/ml Doxycycline hyclate and 0.1x ROCK inhibitor (RevitaCell Supplement). 72 hrs after Doxycycline induction (defined as day 0 iNeurons), cultures were switched to cortical neuron culture media consisting of BrainPhys (05791; STEMCELL Technologies), 10 μg/mL BDNF (450-02; PeproTech), 10 μg/mL NT-3 (450-03; PeproTech), 2% B-27 (A1486701; Gibco), 1 μg/mL Laminin (23-017-015; Gibco) and supplemented with 0.5 μg/ml Doxycycline hyclate. iNeurons were cultured up to 21 days as adherent monolayers at 37°C in a humidified incubator with 5% CO_2_ with half medium changes every 2-3 days in doxycycline-free cortical neuron culture medium.

### iNeuron transduction

Lentiviral shRNA expression plasmids were from Sigma: MISSION® TRC2 pLKO.5-puro Non-Mammalian shRNA for control, TRCN0000263599 for ATAT1. Lentivirus was concentrated with Lenti-X Concentrator (Takara Bio 631231) in miliQ and treated cells with diluted 1:2000 in cortical media. 24 hours after the transduction, medium was replaced with fresh cortical neuron culture media.

### Brightfield imaging for cell morphology

Brightfield images were acquired using an EVOS M5000 inverted microscope (Thermo Fisher Scientific) equipped with a transmitted-light illumination module using a 20x objective.

### Western blotting

Cells were lysed on ice in RIPA buffer (50 mM Tris, pH 8, 150 mM NaCl, 0.5% deoxycholate, 0.1% SDS, 1% NP-40 supplemented with 15 mM sodium pyrophosphate, 50 mM sodium fluoride, 40 mM β-glycophosphate, 1 mM sodium vanadate, 100 mM phenylmethylsulfonylfluoride, 1 μg/mL leupeptin, and 5 μg/mL aprotinin) supplemented with 1:100 Protease Inhibitor Cocktail (Sigma; P8340-5mL), 1:100 Phosphatase Inhibitor Cocktail 2 (Sigma; P5726), 1:100 Phosphatase Inhibitor Cocktail 3 (Sigma; P0044). Adherent samples were scraped, collected, and centrifuged at 13,000 xg for 5 min at 4°C to isolate the post-nuclear supernatant. Concentration of supernatant was measured by Pierce BCA Protein Assay Kits (Thermo Scientific; #23225), and the supernatant were then boiled with 6x Laemmli buffer containing β-mercaptoethanol and denatured for 10 min at 95°C. Equal protein amounts were loaded into each well of 10% SDS-PAGE and transferred onto 0.2 μm nitrocellulose membrane (Bio-Rad 162-0112) by wet transfer 1 h at 100 V for immunoblotting. Membranes were blocked in 5% milk in Tris-buffered saline (TBS) for 30 min at room temperature and then incubated with primary antibodies diluted in 5% milk in TBST overnight at 4°C. The next day, membranes were washed three times 5 mins with TBST, incubated with secondary antibodies diluted in 5% milk in TBST 1 h at room temperature, washed three times 5 mins with TBST, and imaged using an Odyssey CLx (LI-COR Biosciences). Western blots were quantified using LI-COR Image Studio. Background was subtracted for each quantified band.

### Microtubule PTM distribution assay

Cells were fixed with ice-cold 100% methanol for 10 mins at −20°C, then washed with DPBS 3 times. Cells were blocked using a blocking solution of 4% bovine serum albumin (BSA) in DPBS for 1 hour. Primary antibodies against ankyrin G (mouse, 1:100) and either acetylated tubulin (rabbit, 1:400) or polyglutamylated tubulin (rabbit, 1:300) were diluted in blocking solution and incubated with cells for overnight at 4°C. After primary antibody incubation, cells were washed with DPBS in three intervals of 5 minutes. Secondary antibodies, anti-rabbit 568 (1:500) and anti-mouse 647 (1:500), were diluted in blocking solution at room temperature for 1 hour while covered from light. After incubation, the secondary antibodies were washed out using three 5-minute DPBS washes. FlexAble CoraLite plus 488 antibody labeling kit for mouse IgG (Proteintech KFA021) was used in 4% BSA in DPBS to label DM1A primary antibodies (anti-alpha-tubulin, mouse) (1:500) for overnight at 4°C. The FlexAble solution was washed out using three 5-minute DPBS washes before DAPI was added in DPBS (1:1000) and incubated at room temperature for 10 mins. The DAPI solution was replaced with DPBS, and the samples were imaged using a Zeiss laser scanning microscope (LSM) 800 with a 63x 1.4 NA oil objective. Airyscan Z-stacks images were collected and processed using ZEN 2.3 Airyscan 3D Processing software. Sum projections of the central region were generated with Fiji-ImageJ for quantification in CellProfiler. Images were evaluated for adequate presence of the AnkG pre-axonal segment marker to confirm that they were properly differentiating into cortical neurons before quantification.

### Multispectral imaging of organelles

Day 6 iNeurons were co-transfected with six piggyBac organelle reporter plasmids together with a piggyBac transposase plasmid using Lipofectamine 2000 (11668027; Thermo Fisher Scientific) following a modified version of the manufacturer’s protocol. For each well of an 8-well chamber, plasmid DNA (120ng EIF1α-Transposase, 120 ng LAMP1::mTurquoise for lysosome, 120 ng Cox8::eGFP for mitochondria, 120 ng mApple::Sec61B for ER, 60ng SiT::oxVenus for Golgi, 25ng mOrange:: SKL for peroxisome, and 60 ng mTagBFP2::NLS for nucleus) were diluted in 25 µL Neurobasal media (12348017; Gibco). In parallel, 1 µL Lipofectamine 2000 was diluted in 25 µL neurobasal media and incubated for 5 min at room temperature. The diluted DNA and Lipofectamine 2000 solutions were then combined, properly mixed and incubated for 15 min at room temperature. Prior to transfection, cultures were washed and equilibrated with neurobasal medium. After a 45-min incubation with the transfection mixture at 37°C and 5% CO_2_, cultures were washed once and replaced with BrainPhys Imaging Optimized Medium (05796; STEMCELL Technologies) supplemented with 2% NeuroCult SM1 (05711; STEMCELL Technologies), 10 μg/mL BDNF (450-02; PeproTech), 10 μg/mL NT-3 (450-03; PeproTech), and 1 μg/mL Laminin (23-017-015; Gibco). On day 7 of iNeuron, 15 min prior to live-cell imaging, 1:2000 CellMask Deep Red (C10046; Invitrogen) was added to label the plasma membrane and 0.5 µM Lipi-Blue (LD01; DOJINDO Laboratories) was added to label lipid droplets. For single-label controls, each plasmid or dye was applied individually using the same DNA-to-Lipofectamine ratios and dye concentrations as in multi-labeled conditions.

### Multispectral imaging

Multispectral imaging was carried out as previously described^41,43,59^. Images were acquired using a Zeiss LSM 880 laser scanning confocal microscope on an inverted Axio Observer Z1 microscope and equipped with a 34-channel GaAsP spectral detector and a live-cell incubation chamber (Carl Zeiss Microscopy). 405 nm (0.5%), 458 nm (3%), 514 nm (0.4%), 561 nm (0.3%), and 633 nm (0.2%) lasers were used simultaneously to excite cells labeled with one or multiple organelle markers. Emitted fluorescence was collected in lambda mode using a linear array of 34 photomultiplier tubes (PMTs) with 9 nm spectral bins spanning 410-695 nm, resulting in 32 intensity channels per image. Z stack images were acquired by a plan-Apo 63x/1.4 NA oil-immersion objective with an XYZ voxel size of 0.07 µm × 0.07 µm × 0.39 µm. Z-stacks were acquired at a single time point with a pixel density of 796 × 796 per slice, and each Z-slice was captured in 1.97s. The number of Z-slices per image was determined on a cell-by-cell basis to capture the entire cell volume from basal to apical surfaces. All imaging was performed live and maintained at 37°C and 5% CO_2_ during imaging.

### Multispectral image processing

Using the linear unmixing algorithm from ZEN Black software version 2.3 (Carl Zeiss Microscopy), spectral Z stack images were first processed into 8-channel images. Control images labeled with a single fluorophore corresponding to one of the eight organelle markers were acquired as described above and used to generate reference spectra. For each fluorophore, a representative region of fluorescence intensity was selected to define its reference spectrum. All reference spectra were then used as inputs and applied to all images across the full biological replicate dataset for linear unmixing, resulting in an eight-channel image in which each channel represents fluorescence intensity from a single fluorophore.

Linear unmixed images were subsequently deconvolved using Huygens Essential software version 23.04 (Scientific Volume Imaging, The Netherlands, http://svi.nl). Excitation and emission maxima (Ex/Em max) for each fluorophore were estimated from the lasers used during imaging and the corresponding reference spectra and applied for batch deconvolution of eight intensity channels using the Workflow Processor module.

### Antibodies and live cell imaging reagents

Antibodies used were: rabbit Monoclonal against acetyl-α-tubulin (Lys40) (D20G3) (Cell signaling #5335; 1:1000 for immunofluorescence staining (IF), 1:5000 for western blotting (WB)), mouse monoclonal against acetyl-α-tubulin (Lys40) (clone 6-11B-1, Sigma #T7451, 1:1000 IF, 1:5000 WB), mouse against α-tubulin DM1A (Abcam #ab7291; 1:1000 IF, 1:10000 WB), rabbit monoclonal against β-III tubulin (Abcam ab18207; 1:1000 WB), mouse monoclonal against actin (SPM161) (Santa Cruz sc-56459; 1:5000 WB), rabbit polyclonal against detyrosinated tubulin (Merck AB3210; 1:250 IF, 1:5000 WB), rabbit polyclonal against ATAT1 (Proteintech 28828-1-AP; 1:3000 WB), mouse monoclonal against ankyrin-G (N106/36) (antibodiesinc SKU: 75-146; 1:500 IF), rabbit polyclonal anti-Polyglutamate chain (polyE), pAb (IN105)(AdipoGen Life Sciences AG-25B-0030-C050; 1:300 IF, 1:5000 WB).

### STED imaging

12mm dia. #1.5H thick precision coverslips (Neuvitro corporation, SKU: GG-12-15H) were sonicated with 70% ethanol for 30mins three times and then washed with autoclaved miliQ once and placed in 24 well Fisherbrand™ Surface Treated SterileTissue Culture Plates (FB012929) UV to dry for more than 15 mins. Coated the coverslips with 0.1mg/mL Poly-L-ornithine hydrobromide (Sigma; #P3655) in Borate buffer (10 mM Boric acid (Sigma B6768), 2.5 mM Sodium tetraborate (Sigma 221732), 7.5 mM Sodium chloride (Sigma S7653), 0.1M Sodium hydroxide (Sigma 71463)) overnight. Then washed with autoclaved MiliQ water 3 times then coated with 10ug/mL Laminin (Gibco™ Laminin Mouse Protein, Natural, 1mg; 23-017-015) in cold DPBS for 2 hours at 37°C. Then cultured iNeuron on PLO/Laminin coated coverslips till day 7 or day 14. iNeurons were fixed with warm 4% paraformaldehyde (PFA) (Electron Microscopy Sciences #15710) in PHEM buffer (60 mM PIPES, 25 mM HEPES, 10 mM EGTA, 2 mM MgSO4, and 0.12 M sucrose) at 37°C for 2 min, followed by 3 min ice cold 100% Methanol incubation at −20°C. Fixed neurons were washed three times in 1xDPBS and permeabilized for 10 min in 500 µl/slide of 0.2% Triton X-100 in DPBS. Samples were further incubated in 100 µl/slide of blocking buffer 4% Donkey serum (Sigma; S30-M) in 0.1% triton in DPBS for at least 45 min at room temperature. Then cells were co-labeled with primary antibodies diluted in blocking buffer for 4°C overnight. (Primary antibodies: mouse anti-AcetyTub 1:300 (Sigma T7451); chicken anti-α-tubulin monoclonal recombinant IgY 1:300 (SYSY antibodies; #302 209); goat anti-Calnexin antibody-C-terminal 1:200 (abcam; ab219644); Rabbit anti-LAMP1 (D2D11) XP mAb 1:500 (Cellsignaling; #9091) Cells were washed with 5mins 1xDPBS three times then followed by species specific secondary antibodies in blocking buffer for 1 hour room temperature, donkey anti-mouse IgG (H+L), Highly Cross-Adsorbed, CF640R 1:200 (Biotium; #20177-1); donkey anti-chicken IgY (H+L) Highly Cross Adsorbed Alexa Fluor™ 594 1:200 (Invitrogen; A78951); donkey anti-goat IgG (H+L), Highly Cross-Adsorbed,CF680R 1:200 (Biotium; #20196-1); donkey anti-rabbit IgG (H+L) Highly Cross-Adsorbed Alexa Fluor™ Plus 488 1:300 (Invitrogen; A32790). Cells were embedded in ProLong™ Glass Antifade Mountant for 48 hours before imaging. Stained samples were acquired on a Leica STELLARIS 8 FALCON STED on an inverted Leica DMI8 motorized microscope (Leica microsystems) with 775nm depletion laser, White Light Lasers (WLL) for excitation, and HyD detectors using HC PL APO 100×/1.40 Oil objective. Images were acquired sequentially in the order 640 nm, 680 nm, 594 nm then 488 nm. For three-color 3D STED imaging of CF640R, CF680R, and Alexa Fluor 594, excitation was performed using 637 nm, 685 nm, and 594 nm laser, respectively. A 775 nm depletion laser was applied at 50%, 50%, and 80% power, respectively, with three-line accumulations for each channel. Alexa Fluor 488 was acquired in confocal mode using 499 nm excitation with the same objective, optical settings, and zoom as used for STED imaging, but without signal averaging or accumulation. Emission detection windows were 653-681 nm, 698-747 nm, 600-637 nm, and 509-597 nm, respectively. All images were acquired as four channels, three Z-stack slices, imaged sequentially from the basal to apical direction with an imaging format of 352 × 352 pixels and an optical zoom of 8, resulting in a voxel size of 40 nm x 40 nm x 50 nm (x/y/z). Scanning was performed in unidirectional mode at a scan speed of 600 Hz. The Z-stack images were subjected to deconvolution using Huygens Essential software version 24.10 (Scientific Volume Imaging, The Netherlands, http://svi.nl) with CMLE (classic maximum likelihood estimation) algorithm with parameters of −100 acuity and 8-11 SNR (Signal-to-Noise Ratio).

### Live-cell lysosome dynamics and functional staining

To measure lysosomal tubulation dynamics, cells were labeled using the Magic Red Fluorescent Cathepsin B Assay Kit (ImmunoChemistry Technologies, 937), SiR-Lysosome Kit (Cytoskeleton, CY-SC012), and LysoTracker Green DND-26 (Invitrogen, L7526). Cells were incubated with either the Magic Red Kit (1:500) or SiR-Lysosome Kit (1:1000) for 30 min at 37 °C, washed once with complete medium, followed by incubation with LysoTracker Green (50 nM) for 15 min at 37 °C prior to imaging. Following staining, time-lapse imaging was performed on microscope with a 1s frame interval for 3 min. Lysosomal acidity was measured using the Lysosomal Acidic pH Detection Kit-Green/Red (Dojindo, L266-10) according to the manufacturer’s instructions. Cells were washed once with complete medium then incubated with LysoPrime Green (1:2000) for 30 min at 37 °C, washed twice with complete medium, followed by incubation with pHLys Red (1:1000) for 30 min at 37°C. Cells were washed twice with complete medium prior to imaging. Autophagy assay used an Autophagic Flux Assay Kit (Dojindo, A562-10) according to the manufacturer’s instructions. Cells were washed once with complete medium then incubated with DALGreen (1:1000) and DAPRed (1:500) for 30 min at 37 °C, then washed twice with complete medium prior to imaging. All lysosome dynamics and functional assays were imaged on a Zeiss laser scanning microscope (LSM) 800 with a 63x 1.4 NA oil objective.

### Quantification and statistical analysis

#### 3D multispectral image segmentation

Instance organelle segmentations were generated from independent intensity channels in the deconvolved Z stack images. Detailed segmentation methods are described in the image analysis pipeline in our Python-based segmentation and analysis package, infer-subc (https://github.com/SCohenLab/infer-subc/tree/v2.0.0b1). Lysosome and mitochondria object masks were the only organelles to be processed using the optional declumping step that exists at the end of each organelle segmentation workflow. Whole cell masks (CellMask) and nuclear masks were generated using segmentation workflow 1.1c, followed by manual refinement in Napari based on plasma membrane and ER signals. CellMasks were subsequently processed using a dedicated workflow to segment the soma and neurites into separate soma and neurite masks (segmentation workflow 1.8).

#### 3D multispectral organelle quantification

Quantitative analysis of 3D organelle organization was performed as previously described^43^ and detailed in infer-subc (https://github.com/SCohenLab/infer-subc/tree/v2.0.0b1). Following segmentation, individual organelle objects and inter-organelle interactions were quantified. Organelle interactions were defined as regions of overlap between organelle objects and were assessed for all combinations among the six organelles. For each organelle and organelle interaction, features including morphology (amount, volume, and shape) and spatial distribution were extracted. Analyses were performed from the full cellular volume captured in each imagewhich was further divided into soma and neurite sub-regions. All metrics were quantified per-subregion using the infer_subc analysis pipeline (”organelle-signature-analysis.ipynb”). After per-object measurements were collected, they were summarized per-subregion, resulting in a total of 5439 organelle metrics within the soma subregion or 3035 metrics from the neurites subregion analysis (distribution analysis not included).

The metrics for the soma and neurites analyses are featured separately throughout this paper. To reduce redundancy and improve power for statistical comparisons, the 5439 soma-specific organelle metrics were narrowed to a curated set of 449 metrics, and the 3035 neurite-specific metrics to a curated set of 131 metrics, which were used to identify major differences between conditions.

### Microtubule PTM distribution analysis

The nucleus and soma regions were manually segmented and saved as a binary mask for region of analysis using CellProfiler. Next, the analysis was performed using an automated pipeline in CellProfiler. Briefly, the intensities of both the total α-tubulin (DM1A) and the PTM tubulin (Acetyl or PolyE tubulin) were rescaled to a ratio of 0-1. Then nuclear and soma masks were applied to define the cytoplasmic region (soma excluding the nucleus), which was further divided into three radially equidistant regions, the perinuclear, the central, and the peripheral regions according to distance from the nucleus for spatial quantification. The average rescaled intensity of the PTM tubulin in each region was then divided by the rescaled intensity of the total α-tubulin to give the radial mean intensity fraction.

### STED image analysis

Deconvolved four-channel STED/confocal 3D Z stack images (three Z-slices) were sum-projected along the Z-axis in Fiji-ImageJ then imported into CellProfiler (v 4.2.6) for segmentation and analysis. Briefly, the cytoplasmic soma area excluding the nucleus was manually selected and saved as a binary mask for region of analysis. Each channel was first intensity-rescaled and then segmented. ER were segmented as a single merged object per cell. The tubeness of total α-tubulin and acetylated tubulin signals were enhanced by 2.0 smoothing scales prior to segmentation. After generating object masks for each channel, the total object-occupied area, object number, and mean object area were quantified for ER, lysosomes, lysosome-ER contacts, acetylated tubulin, and non-acetylated tubulin. Lysosome-ER contact sites were quantified by measuring the overlap between lysosomes and ER masks. Non-acetylated tubulin regions were defined by subtracting the acetylated tubulin mask from the total α-tubulin mask. Lysosome-ER contacts localized to acetylated tubulin or non-acetylated tubulin were quantified by measuring overlap between Lysosome-ER contact masks and acetylated tubulin or non-acetylated tubulin masks, respectively. All conditions were quantified using an identical CellProfiler pipeline with fixed parameters, applied in batch mode.

### Tubular lysosome ratio measurements

Timelapse images with LysoTracker Green and Magic Red labeling were deconvolved using Huygens Essential software version 24.10 (Scientific Volume Imaging, The Netherlands, http://svi.nl). Next, lysosome tubulation events were marked and quantified based on LysoTracker Green channel using the Cell Counter plugin in Fiji. Lysosome tubulation was defined as protrusions from lysosomal bodies resulting in retraction, stable tubules, or fission of the protrusion from the body. Tubulation events were identified and classified in one of the three above categories throughout the 3 mins time course. Mature lysosomes were determined based on the presence of Magic Red signals. Magic Red-positive lysosomes were segmented using the Yen auto thresholding method with the stack histogram option applied in Fiji. For each identified lysosomal tubulation event, the lysosomal body was classified based on whether the lysosomal body overlapped with segmented mature lysosome regions at any time point during the 3 min time course.

### Lysosomal acidity and autophagy analysis

Lysosomal acidity and autophagy based on live cell dual color sensors were analyzed using a semiautomated image analysis pipeline in Fiji. Cytoplasmic regions excluding the nucleus were first manually segmented and saved as regions of interest (ROIs). For intensity-based measurements, background subtraction was performed on each channel using a 50-pixel rolling radius, and mean fluorescence intensities within cytoplasmic ROIs were measured for each channel. The fluorescence intensity ratio between channels was calculated. For acidic lysosome area analysis, cytoplasmic ROIs were imported into a custom automated Fiji pipeline. Images were first smoothed using a Gaussian blur (σ = 2.0) and non-cytoplasmic regions were removed using the “Clear Outside” function.

Lysosomes were segmented using the Bernsen auto local threshold with a radius of 15 pixels, followed by watershed separation. Segmented objects larger than 15 pixels were retained and the number of objects, total area, and individual object area were quantified using Analyze Particles.

### Statistics

All experiments were performed with a minimum of three independent biological replicates unless otherwise specified. Each replicate was from a separate cell culture. The number of replicates and statistical tests are indicated in the figure legends. For comparisons between two groups, T-tests were used. For multiple-group comparisons, one-way ANOVA followed by appropriate post hoc tests (specified in the figure legends) was performed. For microscopy experiments, fields of view were selected at random, but image acquisition and analysis were not performed blinded to experimental conditions. For multispectral imaging datasets, a nonparametric Mann-Whitney U test and statistical significance was determined by a 10% false discovery rate. All statistical analyses, plots and heatmaps were conducted using GraphPad Prism 10 (RRID:SCR_002798) Windows, GraphPad Software, Boston, Massachusetts USA, www.graphpad.com

## Data availability

The data generated in this study are available from the corresponding author upon reasonable request.

## Acknowledgements

We thank Dr. Bill Skarnes (The Jackson Laboratory) for generously providing the KOLF2.1J cell line. We thank Dr. Michael Ward (NINDS) for kindly sharing piggyBac vectors. We thank Wendy Salmon (Hooker Imaging Core) for expert imaging advice. Multispectral microscopy (Zeiss 880 Confocal) was performed at the UNC Hooker Imaging Core Facility, supported in part by P30 CA016086 Cancer Center Core support grant to the UNC Lineberger Comprehensive Cancer Center. The Leica STED instrument at the UNC Hooker Imaging Core Facility is supported by National Institutes of Health grant 1S10OD030300. This work was supported by the National Institutes of Health under award R35GM133460 (S.C.), by Chan Zuckerberg Initiative award A23-0264-001 (S.C.) and by the Ministry of Education, Taiwan, under the Government Scholarship to Study Abroad (GSSA) program (C-H.H.). We thank Shannon Rhoads, Ngudiankama Rene Mfulama and Zachary Coman for assistance with the 3D organelle analysis pipeline (infer-subc).

## Contributions

C-H.H. and S.C. conceived the study, obtained funding, and wrote the paper. C-H.H. and A.J.K. performed the experiments and analyzed the data. M.C.Z. contributed key research materials and data analysis tools. S.C. supervised the project.

## Ethics declarations

The authors declare no competing interests.

